# A CRISPR endonuclease gene drive reveals two distinct mechanisms of inheritance bias

**DOI:** 10.1101/2020.12.15.421271

**Authors:** Sebald A. N. Verkuijl, Estela Gonzalez, Ming Li, Joshua Ang, Nikolay P. Kandul, Michelle A. E. Anderson, Omar S. Akbari, Michael B. Bonsall, Luke Alphey

**Author notes:** these authors contributed equally to this work (Sebald A. N. Verkuijl, Estela Gonzalez).

## Abstract

RNA guided CRISPR gene drives have shown the capability of biasing transgene inheritance in multiple species. Among these, homing endonuclease drives are the most developed. In this study, we report the functioning of *sds3, bgcn*, and *nup50* expressed Cas9 in an *Aedes aegypti* homing split drive system targeting the *white* gene. We report their inheritance biasing capability, propensity for maternal deposition, and zygotic/somatic expression. Additionally, by making use of the tight linkage of *white* to the sex-determining locus, we were able to elucidate mechanisms of inheritance bias. We find inheritance bias through homing in double heterozygous males, but find that a previous report of the same drive occurred through meiotic drive. We propose that other previously reported ‘homing’ design gene drives may in fact bias their inheritance through other mechanisms with important implications for gene drive design.

## Introduction

Genetic modification of wild populations has been proposed as a means of addressing some of the world’s most pressing public health challenges and may be achieved through gene drive. Gene drive is the ability of a genetic element to bias its own inheritance, which allows it to spread a genetic change through a population without necessarily conferring a fitness benefit (’selfish DNA’)^1^. There are many examples of this phenomenon in nature, acting through many different mechanisms^2^. Some types of gene drive rely on the action of sequence-specific DNA nucleases (enzymes that create DNA breaks). Double-stranded DNA breaks are a common occurrence in cells, and a range of mechanisms exists to repair the DNA damage. Correspondingly, different nuclease-based gene drives can potentially bias their inheritance through different mechanisms. The development of synthetic nuclease gene drives received much attention^3,4^ following the discovery of programmable nucleases in CRISPR systems^5^. Particularly the development of gene drives based on ‘homing’ (homing drives) and drives that cause the loss of non-drive bearing gametes or offspring (here referred to as meiotic drive).

Generally, in diploid organisms, one chromosome of each homologous pair is contributed by each parent, and each allele has a 50% chance of being passed along to a given progeny. Synthetic homing and meiotic endonuclease gene drives both rely on selectively creating double-strand DNA breaks on the non-drive-bearing homolog when the drive is present on one of the pair but not the other (heterozygous). Through different mechanisms, this results in an inheritance bias of an allele or genomic region and for meiotic drive potentially the whole chromosome. Meiotic endonuclease drives lower the inheritance of the competing chromosome within a pair by damaging it such that gametes carrying the non-drive chromosome are eliminated during gametogenesis or, in some cases, produce non-viable offspring. This includes the disruption of specific essential genes in toxin-antidote meiotic drives^6–8^, or through more structural damage such as with chromosome ‘shredder’ meiotic drives^9,10^. Natural sex-linked meiotic drive systems have been reported in *Aedes* and *Culex* mosquitoes^11,12^. Synthetic shredder endonuclease meiotic drives have generally sought to exploit naturally present large-scale, potentially repeating, sequence differences between chromosome pairs^9,10^. In contrast to meiotic drives, for homing drives sequence homology between the drive element and the target on the paired chromosome is essential. Homing drives bias their inheritance by creating a DNA break on the ‘recipient’ homologous chromosome corresponding to where the genetic material of the homing drive is located on the ‘donor’ homologous chromosome. If the coding sequence for the drive is then identified as missing from the cut chromosome, the DNA sequences of the drive and associated sequences can be copied over during repair of the DNA break (’homology-dependent repair’).

For most reports of synthetic homing drives, the method of quantifying inheritance bias (phenotypic scoring of progeny carrying a drive linked genetic marker) cannot differentiate between the underlying inheritance biasing mechanism^13–34^. The large differences in ‘design rules’ that have emerged between synthetic meiotic and homing endonuclease systems may have contributed to the expectation that, for any given neutral target drive, there should be little overlap in the mechanism. This is supported by a (small) subset of publications that have used marked chromosomes^35–37^, especially pre-CRISPR^38–41^, which allowed the homing and meiotic drive mechanisms to be differentiated. These studies did not report observing meiotic drive. However, we noted evidence for meiotic drive in male *Aedes aegypti* with a homing CRISPR gene drive design recently reported in Li et al.^36^.

Li et al. tested the inheritance biasing ability of a set of ‘homing’ split-drive systems comprising a gRNA expressing element inserted into the *white* gene (*w*^GDe^) and one of four second-site transgene insertions expressing Cas9 under the control of various promoters from genes expressed in the mosquito germline. They demonstrated that *w*^GDe^ is tightly linked to the sex-determining region of *A. aegypti*, a region on chromosome one with a dominant male-determining allele *M* such that males are *M*/*m* and females *m*/*m*. This allows the sex of progeny to function as a chromosomal marker in the progeny of male drive carriers. While three of the *Cas9* regulatory regions resulted in drive activity in females, only ‘*nup50*’ expressing Cas9 resulted in a statistically significant increased inheritance of the drive from male drive parents. We reanalysed the results of Li et al. for *nup50* males taking into account the sex linkage and found that the observed inheritance bias in double heterozygous males proceeded exclusively through meiotic drive.

We set out to test the hypothesis that the apparent meiotic drive observed with the *nup50* expression pattern is a more general phenomenon and also occurs with other *A. aegypti* gene drives that show activity in males. In collaboration with the original authors, we repeated the *nup50*-*Cas9* crosses and performed crosses with Cas9 expression under the control of putative transcriptional regulatory regions from two additional *A. aegypti* germline genes. The first, suppressor of defective silencing 3 (*sds3*) has been shown, by dsRNA-induced knockdown in *Anopheles gambiae*, to be required for normal development of the ovarian follicles and testis, with no other obvious defects^42^. The second, benign gonial cell neoplasm protein (*bgcn*) is involved in regulating and promoting gametogenesis in both sexes^43^ and has been described in the context of gene drive in *Drosophila melanogaster* with I-*Sce*I^38^ and *A. aegypti*^44^.

For each Cas9 expressing line, we report the degree of inheritance bias of the *w*^GDe^ element for both sexes and, in males, the mechanism of inheritance bias. In addition, by scoring somatic eye phenotypes, we also find strong evidence of zygotic/somatic expression, maternal deposition and unexpectedly a currently unexplained effect of the *Cas9* carrying grandparent’s sex on *w*^GDe^ grand-offspring phenotypes.

## Methods

### DNA constructs

The *sds3*-*Cas9* construct was produced by making several alterations to plasmids provided by Omar Akbari^45^. The *sds3* construct contains, within piggyBac terminal sequences, an insect codon optimised *Cas9* followed by a T2A self-cleaving peptide and EGFP. To improve visibility of the fluorescent marker, the initial OpIE2-DsRED cassette was replaced with PUb-mCherry-SV40. To reproduce the germline-specific expression pattern of *sds3*, the Cas9:EGFP coding sequence is preceded and followed by the non-coding sequences flanking the endogenous *sds3* gene’s open reading frame. The 3’ UTR is followed by an additional P10 3’UTR. To determine the 5’ and 3’ UTR of the *sds3* gene 5’ and 3’ Rapid amplification of cDNA ends (RACE) was performed using the SMARTer® RACE 5’/3’ Kit (Takara Bio) on RNA isolated using Trizol (Life Technologies) from female and male, 5-7 day post eclosion WT adult mosquitoes. RACE PCRs were performed using gene-specific primers: 5’-TGTGCTGTTCGTATGGTTCCGGATGG-3’ then nested primer 5’-TCGTCCAGCAAAAGAACCAACTGCCCAG-3’ for 5’ RACE and for 3’RACE 5’-ACGTCGACCTAATGAACCGCTTCCG-3’ and nested primer 5’-GGGCAGTTGGTTCTTTTGCTGGACG-3’, amplicons were cloned using the CloneJET PCR Cloning Kit (Thermo Scientific) and Sanger sequenced. In total, 1959 bp upstream of the translational start was amplified in order to include the most significant promoter elements (including 202 bp of 5’UTR). The 5’ and 3’ sequences were amplified from WT adult gDNA extracted using the NucleoSpin Tissue Kit (Machery-Nagel) by PCR using Phusion High Fidelity PCR Master Mix (NEB) and primers 5’-tttt<u>gcggccgc</u>TCTGTTTGAATATGTTTCCGAGAA −3’ and 5’-tttt<u>ctcgag</u>TTTCCGCGACAAAAACACAGA-3’ which add restriction enzyme recognition sites *Not*I and *Xho*I, respectively (underlined).The promoter fragment digested with *Not*I/*Xho*I, *Cas9* digested with *Xho*I/*Fse*I, and pBac PUb-mCherry-SV40 digested with *Not*I and *Fse*I were ligated with T4 DNA ligase (NEB) in a three-way ligation. The 307 bp 3’UTR was then amplified from the same WT adult genomic DNA by PCR using Phusion High Fidelity Master Mix (NEB) and primers: 5’-ttttttaattaaGGAAACAAGGATCTCAACTCTCGAGC-3’ and 5’ttttg cgatcgcc<u>ctcgag</u>cTTCTTAGGTACAATTGTAAAACATAGTT-3’ to amplify from the stop-codon until just beyond the transcript end as determined by 3’ RACE. This amplicon was digested and ligated into the *sds3*-*Cas9* intermediate plasmid digested with *Pac*I. See GenBank depository for the sequences and annotations of *sds3*. The *bgcn* construct was created similarly to the *sds3* construct and is described in Anderson et al.^44^. The sequence and insertion site of the gRNA element (3xP3-tdTomato) and *nup50* lines are described in Li et al.^36^. The *bgcn* and *sds3* constructs use a *Cas9* that is insect codon optimised as described in Anderson et al.^44^. The *nup50* line makes use of a human codon optimised *Cas9*^45^.

### Mosquito lines

*A. aegypti* Liverpool strain (WT) was a gift from Jarek Krzywinski. The *nup50*-*Cas9* and white gRNA expressing element (w^U6b-GDe^, hereafter *w*^GDe^) lines provided by Omar Akbari are described in Li et al.^36^. At Pirbright, the *nup50*-*Cas9* line was maintained as a mix of homozygotes and heterozygotes with periodic selective elimination of wildtypes; the *w*^GDe^ element line was reared as homozygous in our facilities. *Cas9* expressing lines generated at the Pirbright facilities were maintained as heterozygotes, usually by crossing transgenic males to WT females.

All mosquito lines were reared in an insectary facility under constant conditions of 28^°^C, 65-75% relative humidity and 12:12 light/dark cycle (1h dawn/1h dusk). Larvae were fed ground TetraMin flake fish food (TetraMin) while adults were provided with 10% sucrose solution *ad libitum*. Defibrinated horse blood (TCS Bioscience) was provided using a Hemotek membrane feeding system (Hemotek Ltd) covered with Parafilm (Bemis).

### Generation of *sds3*-*Cas9* lines

*sds3*-*Cas9* lines were generated by embryonic microinjection of 1hr post-oviposition WT *A. aegypti* with 500 ng/*µ*l piggyBac donor and 300 ng/*µ*l PUb-piggyBac transposase^44,46^ in 1*×* injection buffer^47^ (Table S1). Injection survivors (G0) were reared to adults and females crossed in pools of approximately 20. Male injection survivors were first crossed individually to 5 virgin WT females then pooled into groups of approximately 20 males. At least 5 ovipositions were collected from each pool, hatched reared to late larvae and screened for the presence of the fluorescence marker on a Leica M165FC fluorescence microscope (Leica Microsystems). From the eight *sds3*-*Cas9* positive pools, a single line was established and used for all the crosses reported here.

### Crosses for homing assessment

Male and female adults, homozygous for *w*^GDe^ were crossed with heterozygous mosquitoes of the *Cas9* lines. Their progeny were screened as late larvae under fluorescence using an MZ165FC microscope. Eye phenotype was also evaluated. Double heterozygous mosquitoes carrying both transgenes were then crossed to WT mosquitoes. Inheritance of the transgenes as well as eye phenotype, was again assessed under a fluorescence microscope. For the *nup50*-Cas9, double heterozygous females from each cross were allowed to lay eggs individually. For *bgcn*-*Cas9* and *sds3*-*Cas9* at least 20 double heterozygotes were crossed separately by sex such that double heterozygotes were always crossed to WT of the opposite sex. The exact number and phenotype of the progeny of each cross are shown in Table S3-S4. The individual cross data for *nup50*-*Cas9* are shown in Table S7-S10. In some cases, F_1_ double heterozygotes produced from the same cross presented with a different fluorescent marker or eye pigment phenotypes. In each case, these were noted in the cross tables, and examples of the phenotypes are shown in Fig S1.

### Statistical methods

#### Statistical analysis of w^GDe^ inheritance bias

For each F_1_ sex, the *w*^GDe^ inheritance rate in the absence of a *Cas9* expressing element (Table S6) was used as the baseline inheritance. This was 52% (620/1203) for males and 51% (308/605) for females. These rates were used as the expected outcome in a Fisher’s exact test with the *w*^GDe^ inheritance from F_1_ parents that carried the *w*^GDe^ and one of the *Cas9* expressing elements. A significant difference in *w*^GDe^ inheritance is taken as evidence for drive activity. See Table S11.

#### Statistical analysis of somatic expression and parental deposition

For each *Cas9* line, the fraction of mosaic eyed (ME) or white-eyed (WE) progeny among the F_2_ offspring inheriting *w*^GDe^ but not the *Cas9* (‘+*w*^GDe^;− *Cas9*’) from F_1_ drive males served as a control for the frequency of such phenotypes in the absence of somatic expression or maternal deposition. For somatic expression, the ME/WE fraction of the F_2_ progeny harbouring both the *Cas9* and *w*^GDe^ elements from F_1_ drive males was compared to the control cross using Fisher’s exact test (Table S12). For maternal deposition, the F_2_ progeny harbouring only the *w*^GDe^ element from F_1_ drive females was compared to the control (Table S13).

#### Statistical analysis of the influence of factors on the fraction of mosaic and white-eyed progeny

A generalised linear model with binomial errors was created that included *Cas9* promoter (*sds3, bgcn, nup50*), F_2_ *Cas9* status (+/−), F_2_ sex (♂/♀), F_1_ drive parent sex (♂/♀), and F_0_ *Cas9* carrying grandparent sex (♂/♀). The response variable was the proportion of ME and WE progeny among the all the F_2_ progeny from that cross and F_2_ sex (48 conditions). The analysis was performed in R version 4.0.2 using the glm function. See Table S14.

#### Statistical analysis of homing and meiotic drive

For homing, the background recombination rate (calculated from the male F_1_ +*w*^GDe^;− *Cas9* cross Table S6) is used as the expected outcome in a Fisher’s exact test. For the control cross (in the absence of possible *Cas9* mediated inheritance bias) the *w*^GDe^ allele was provided by the male F_0_ grandparent and therefore *M*-linked in the male F_1_s. In the absence of recombination, all male F_2_s should be *w*^GDe^ positive, and all female F_2_s should be *w*^GDe^ negative. Out of the 1203 progeny scored, we saw 13 (1.08%) recombination events. 2 out of 609 male F_2_s were *w*^GDe^ negative, and 11 out of 581 female F_2_s were *w*^GDe^ positive. For the crosses including a *Cas9* element, a significant increase in recombination rate between the recipient/donor chromosome marker (sex) and the drive element was taken as evidence of homing (Table S15). For meiotic drive, a statistically significant difference in the inheritance of either the recipient or donor chromosome (i.e. F_2_ sex) is taken as evidence for meiotic drive (Table S16). The progeny sex ratio is compared to the sex-ratio in the absence of a *Cas9* expressing element (Table S6).

## Results

To assess the degree and, in males, the mechanism of inheritance bias, we bred transgenic *A. aegypti* mosquitoes to create and analyse a ‘split drive’ arrangement. In this split drive, the *w*^GDe^ allele expresses a gRNA targeting the wildtype *white* gene (*w*^+^) at the site corresponding to where the drive element has been inserted into and disrupts its protein-coding sequence (Fig 1a). To create individuals in which drive can occur, the *w*^GDe^ element is combined with the other component of the split drive, *Cas9*, as a separate transgene expressing Cas9 under the control of regulatory sequences from an endogenous germline-specific gene, either *nup50, bgcn*, or *sds3*. Individual double heterozygotes for *w*^GDe^ and *Cas9* were generated in two ways: by crossing parental ‘F_0_’ female *w*^GDe^ homozygotes to male *nup50*/*bgcn*/*sds3 Cas9* individuals (Fig 1b Left) or with the reciprocal cross (Fig 1b Right). The double heterozygous offspring (F_1_) were in turn crossed to the Liverpool wildtype strain, and their progeny (F_2_) were collected and scored.

**Figure 1.**
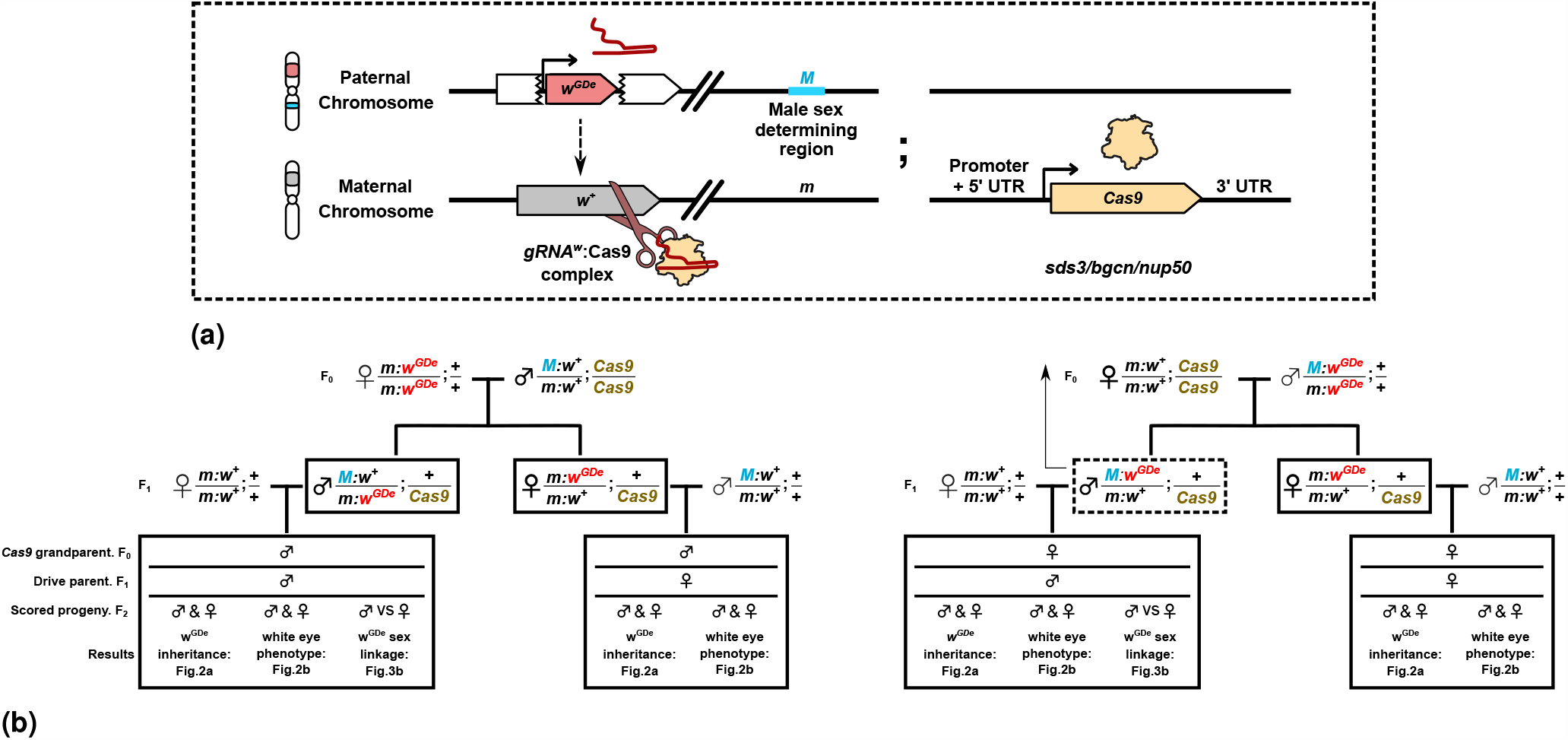
Illustration of split drive design and breeding schemes for the four types of crosses analysed, two for each sex. a. Illustration of *gRNA:Cas9* split-drive system. The gene drive element ^GDe^ is inserted into, and disrupts, the *white* gene which is tightly linked to the sex-determining region (either *M* or *m*). The two loci are separated by ca. 45 Mbp^48,49^. The *w*^GDe^ element illustrated here is *M*-linked on the ‘donor chromosome’. The ‘recipient chromosome’ contains the wildtype *white* gene (*w*^+^), and *m*. Bi-allelic disruption of the *white* gene results in a loss of eye pigmentation. b. (Left) The family tree for double heterozygous male and female parents with paternally contributed *Cas9* and maternally contributed *w*^GDe^. (Right) The family tree for double heterozygous male and female parents with paternally contributed *w*^GDe^ and maternally contributed *Cas9*. The one dashed and three solid boxes indicate the F_1_ genotypes for which germline inheritance bias is measured by scoring their F_2_ progeny. As indicated by the arrow, the F_1_ genotype surrounded by a dashed box is the example illustrated in panel a. A maternally contributed *w*^GDe^ allele (and therefore paternal *Cas9*) results in the drive element being *m*-linked in both male and female F_1_ parents. A paternally contributed *w*^GDe^ allele results in it being *M*-linked in male and *m*-linked in female F_1_ parents. For male F_1_ parents, the *w*^GDe^ allele should be found exclusively in only one sex of their F_2_ progeny, apart from background recombination events, or drive induced recombination (homing).

In Table S2-S4, the progeny genotype and phenotype of the F_2_ progeny are reported for each cross. In Table S11 and Fig 2a the percentage of *w*^GDe^ inheriting F_2_ progeny from double heterozygote F_1_ parents are reported for each Cas9 expressing line. The inheritance rates are split by the sex of the double heterozygous parents, and whether those parents inherited the *Cas9* element from the F_0_ grandmother (maternal) or F_0_ grandfather (paternal). For each condition, Fisher’s Exact tests were performed comparing the *w*^GDe^ inheritance rates to those in the absence of any *Cas9* element for male (52%, 620/1203) or female (51%, 308/605) parents (Table S6).

**Figure 2.**
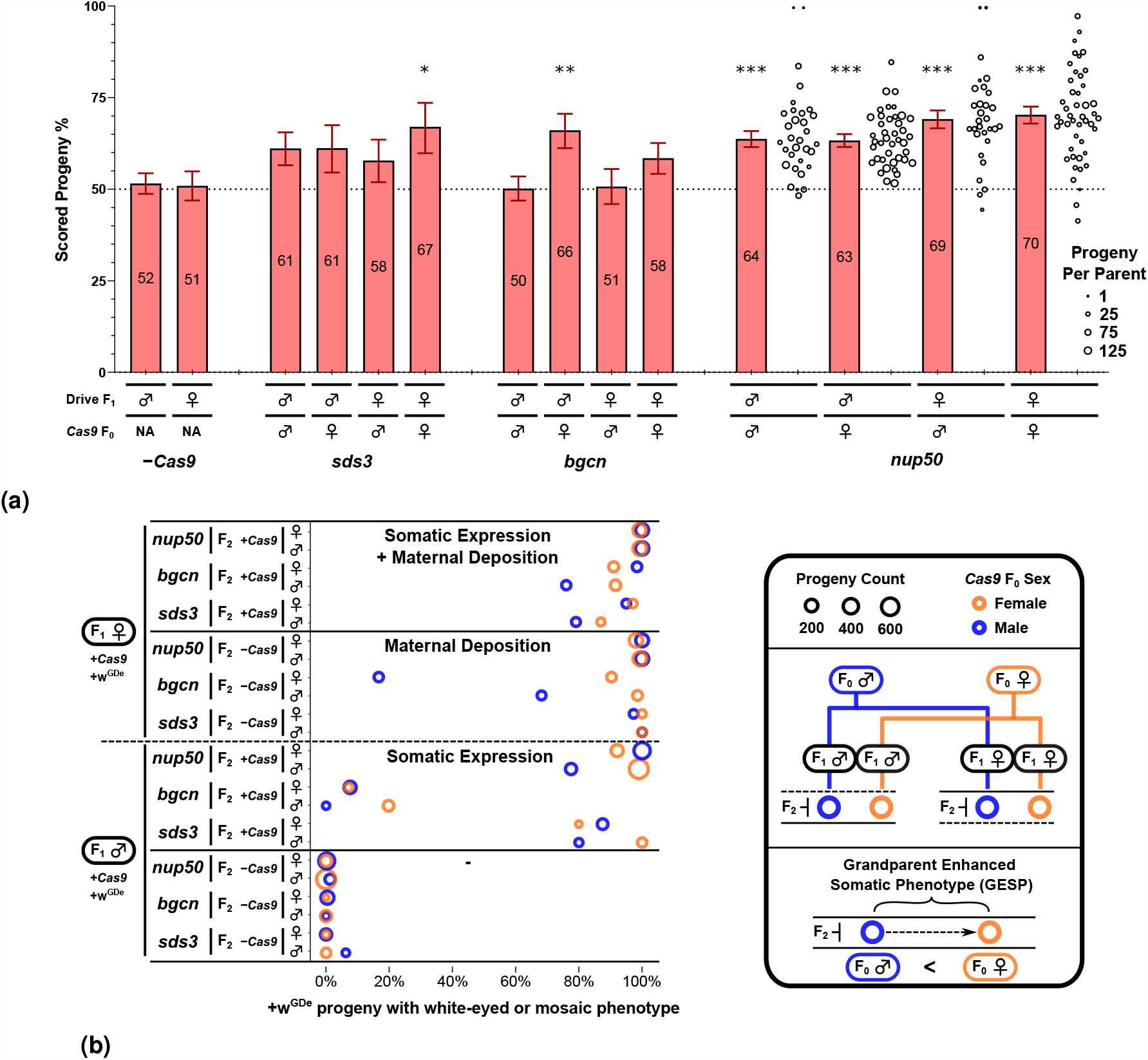
Gene drive element (*w*^GDe^) inheritance and somatic eye phenotype in the progeny of double heterozygote split-drive carriers (’Drive parents’). a Parental germline inheritance bias of *w*^GDe^ when combined with a *nup50, bgcn*, or *sds3*-*Cas9* expressing element. For each of the three *Cas9* regulatory elements, the inheritance rates are reported in columns left-to-right: Male drive F_1_s with paternal *Cas9* contribution, male drive F_1_s with maternal *Cas9* contribution, female drive F_1_s with paternal *Cas9* contribution, and female drive F_1_s with maternal *Cas9* contribution. The horizontal dotted line indicates the expected Mendelian 50% inheritance from heterozygous carriers. For *nup50*, individual crosses were performed, and each circle represents the percentage of *w*^GDe^ positive progeny from an individual parent. Error bars are the Wilson confidence intervals for the binomial proportion calculated for the pooled progeny count, which does not take into account the potential lack of independence due to individual parent ‘batch’ effects. Stars indicate statistical significance as presented in Table S11. b The percentage of +*w*^GDe^ progeny that display a mosaic or total loss of eye pigment phenotype. F_2_ progeny are segregated by the drive carrying F_1_’s sex (♂,♀), the F_2_’s *Cas9* transgene inheritance (− *Cas9*, +*Cas9*), the *Cas9* regulatory sequences (*sds3, bgcn, nup50*), and the F_2_’s sex (♂,♀). The circle size indicates the number of progeny that make up that group, and circle colour indicates if the *Cas9* carrying F_0_ grandparent was male (Blue) or female (Orange). The set of progeny that came from F_1_ drive females are indicated with ‘Maternal Deposition’. The set of progeny that inherited both a *w*^GDe^ allele and *Cas9* element are indicated with ‘Somatic Expression’. Within matched crosses (each row), differences in the white phenotype rate corresponding to the *Cas9* carrying F_0_’s sex are referred to as a grandparent enhanced somatic phenotype. White phenotype rates for − *w*^GDe^ progeny are shown in Fig S2.

Male F_1_ *sds3* double heterozygotes passed along the *w*^GDe^ element to 61% (275/450, p-value: 0.066^ns^) of their progeny with paternal *Cas9* (F_0_: ♂*Cas9*, ♀*w*^GDe^) and 61% (131/214, p-value: 0.157^ns^) of their progeny with maternal *Cas9* (F_0_: ♀*Cas9*, ♂*w*^GDe^). For *sds3* females this was 58% (159/275, p-value: 0.298^ns^) for paternal *Cas9* and 67% (118/176, p-value: 0.050^*^) for maternal *Cas9*. For *bgcn*, only F_1_ drive males with maternal F_0_ *Cas9* had significantly increased *w*^GDe^ propagation rates: 66% (257/389, p-value: 0.010^**^). Similar to biased inheritance rates reported in Li et al., the *nup50* double heterozygous males passed along the *w*^GDe^ element to 64% (1159/1819, p-value: 0.001^***^) of their progeny with paternal *Cas9* and to 63% (1852/2926, p-value: <0.001^***^) of their progeny with maternal *Cas9*. Additionally, for *nup50* drive females this was 69% (952/1377, p-value: <0.001^***^) for paternal *Cas9* and 70% (1055/1501, p-value: <0.001^***^) for maternal *Cas9*.

For *nup50*-*Cas9*, the progeny were collected individually from F_1_ parents (Table S7-S10). As can be seen in Fig 2a, there was considerable variation between the inheritance rate from different parents carrying the same drive, a notable feature that was reported in many other drive papers^18,24,26,35^. Due to this over-dispersal, we cannot reliably determine if there is a statistical difference in the inheritance rate between the different *Cas9* regulatory elements. However, because this over-dispersal is expected only to occur if the drive is functional, our method for determining a difference from the control remains valid, albeit with a potentially inflated false-negative rate.

All progeny were evaluated for eye pigment defects which may result from embryonic or later somatic bi-allelic disruption of the *white* gene by the *w*^GDe^ element and NHEJ mutations. Since the double heterozygote drive carrying parents were crossed to wildtype individuals, each progeny inherited at least one dominant functional *white* allele from the non-drive parent, and, if the *w*^GDe^ element is not inherited, potentially an additional one from the drive parent. Bi-allelic loss of function of the *white* gene must therefore occur through deposition into, or somatic expression in, F_2_ individuals. The progeny from the − *Cas9* control crosses did not present with a white phenotype (Table S6). The eye pigment phenotype for the three Cas9 expressing lines is reported in Fig 2b (+*w*^GDe^) and Fig S2 (− *w*^GDe^).

For the male double heterozygotes *sds3*-*Cas9* crosses, of the F_2_ progeny (♂ and ♀ pooled) that inherited both the *w*^Gde^ and *Cas9* element 86% of presented with a mutant somatic phenotype if the *Cas9* carrying F_0_ was male, and 98% if the *Cas9* carrying F_0_ grandparent was female (F_1_:♂, +*Cas9* in Fig 2b and Table S12). For *bgcn*-*Cas9* this was 7/17%, and for *nup50*-*Cas9* this was 95/98%. However, if only the *w*^GDe^ element was inherited, no cross had more than 1% of the pooled ♂ and ♀ F_2_ progeny present with a somatic phenotype, presumably resulting from the lack of paternal Cas9 transmission through the sperm (F_1_:♂, − *Cas9* in Fig 2b and Table S12). For each cross, this was a significant difference (Table S12) indicating somatic expression, without substantial paternal deposition of Cas9/Cas9:gRNA^w^. In contrast to the <1% rate observed in the progeny of F_1_ drive males, the crosses with female double heterozygotes where only the *w*^GDe^ element was inherited, 40/95% (F_0_:♂/♀) of the *bgcn*, and an astounding 99/100% of the *sds3* and 100/99% of the *nup50* progeny presented with visible somatic phenotypes (F_1_:♀, − *Cas9* in Fig 2b and Table S13). This indicates strong maternal deposition of Cas9/Cas9:gRNA^W^. For each cross, this was a significant difference (Table S13). Maternal Cas9 deposition without substantial paternal deposition has been reported for many other drive systems^13–15,18,24–26,29,30,36, 50–53^.

Surprisingly, in the *w*^GDe^ inheriting progeny we observed a trend where a higher fraction of progeny exhibited a somatic phenotype when the *Cas9* carrying grandparent was female as opposed to male (F_0_:♂ vs F_0_:♀in Fig 2b). Contrasting each male F_0_ *Cas9* carrying grandparent cross with the equivalent cross with a female F_0_ *Cas9* (each row in Fig 2b) showed, for female F_0_ *Cas9*, an average 5.2% (sd:14.4%) percentage point increase in white/mosaic eyed phenotype among +*w*^GDe^ F_2_ progeny. While maternal deposition from a *Cas9* carrying grandparent may increase the number of *w*^GDe^ and NHEJ mutated alleles passed along by the F_1_ parental generation to − *w*^GDe^ progeny (S2), this should not, in contrast to what we observe, influence the phenotype of the progeny that inherit the *w*^GDe^ element (2b). If the *w*^GDe^ element is inherited there is no opportunity to inherit a germline NHEJ mutation that was created due to deposition from the grandparent into the parent. We created a generalised linear model that included *Cas9* promoter, F_2_ *Cas9* status, F_2_ sex, F_1_ drive parent sex, and F_0_ *Cas9* carrying grandparent sex (Table S14). The sex of F_0_ *Cas9* carrying parent had a significant influence on the fraction of white/mosaic eyed +*w*^GDe^ F_2_ progeny. We termed this phenomenon Grandparent Enhanced Somatic Phenotype (GESP). All other factors were significant too, apart from the sex of the F_2_ progeny.

In *A. aegypti*, the *white* gene is tightly linked to the sex-determining locus. This locus comprises two forms, a dominant male-determining allele *M* and a corresponding *m*, such that males are *M*/*m* and females *m*/*m*. While the molecular basis of sex determination in this mosquito is not fully understood, *M* is associated with *Nix*, a gene shown to be involved in sex determination^54^. Analogous to an XY chromosome system, male offspring of an *M*/*m* male always carry the paternal *M* allele and female offspring the paternal *m*, with no such distinction between the two *m* alleles of the mother. For the male parent, if the initial linkage of *w*^GDe^ to either *m* or *M* is known (determined by the sex of the *w*^GDe^ carrying grandparent), the sex of the progeny can be used as an indication of whether an observed inheritance bias is due to new recombination events (homing), or increased inheritance of the original drive carrying chromosome (meiotic drive) (Fig 3a). To this end, we stratified the *w*^GDe^ inheritance by the sex of the F_2_ progeny for each of the male double heterozygous parents (Fig 3b).

**Figure 3.**
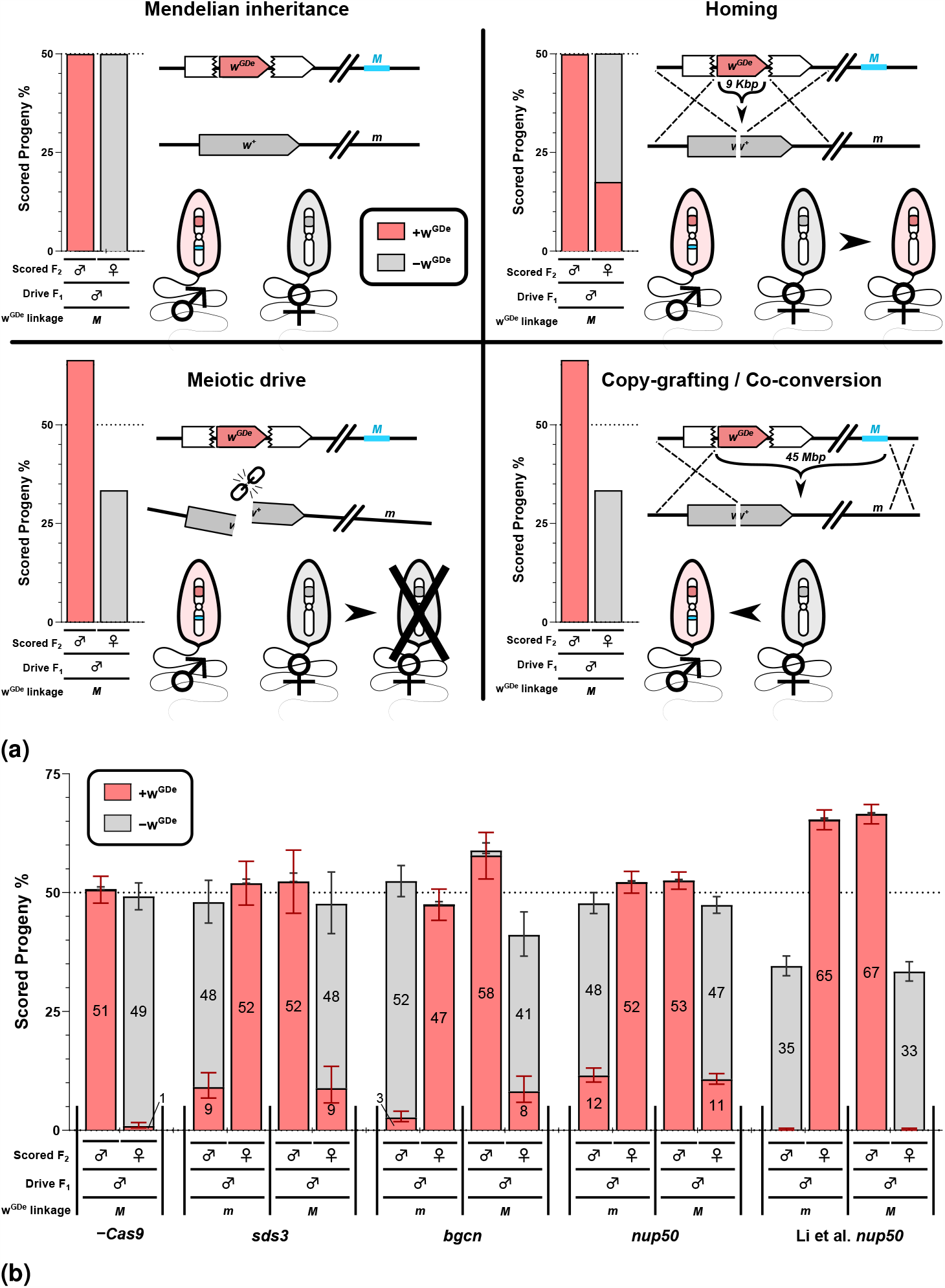
Separating *w*^GDe^ inheritance by F_2_ sex allows different mechanisms of inheritance bias to be distinguished. a Illustration of how homing, meiotic drive and copy-grafting/co-conversion are expected to influence the observed sex-linkage of an *M* linked *w*^GDe^ element in the progeny of male drive double heterozygous parents. b Parental germline inheritance bias of *w*^GDe^ when combined with no *Cas9, nup50, bgcn*, or an *sds3*-*Cas9* expressing element. We included the *nup50* results from Li et al. that use the identical *nup50*-*Cas9* line. For each of the three *Cas9* regulatory elements, the *w*^GDe^ inheritance from male double heterozygotes is reported in pairs of columns segregated by the sex of the F_2_ progeny. In each case, the first pair of columns are the results for when *w*^GDe^ is *m*-linked, and the second pair are the results for when *w*^GDe^ is *M*-linked. Error bars are the Wilson confidence intervals for the binomial proportion calculated for the pooled progeny count. The overlaid numbers are the percentage (cumulative within each column) of the indicated F_2_ sex and *w*^GDe^ status among all progeny from that cross.

The background recombination rate of *w*^GDe^ and sex in the absence of any *Cas9* element was 1.08% (13/1203) (Table S6) and was compared by Fisher’s Exact tests to the recombination rate from *w*^GDe^ *Cas9* male double heterozygotes (Table S15). As reported above, only one cross each of the *sds3* and *bgcn* double heterozygotes showed a significant increase in overall *w*^GDe^ inheritance. However, quantifying conversion with marked chromosomes is much more sensitive than measuring overall *w*^GDe^ inheritance rate.

For the *sds3* double heterozygous males with paternal *Cas9* contribution, 9% of their progeny were *w*^GDe^ positive males.

This indicates that 19% (41/216 p-value: <0.001^***^) of the recipient chromosomes were converted by the combined effect of homing and background recombination. The same was true for maternally contributed *Cas9* where 9% of *w*^GDe^ progeny were female, indicating 19% (19/102 p-value: <0.001^***^) of recipient chromosomes were converted. For *bgcn* males with paternal *Cas9*, we found that 3% of their progeny were *w*^GDe^ positive males which indicates 5% (24/464 p-value: <0.001^***^) of the recipient chromosomes were converted. For *bgcn* with maternally contributed *Cas9*, 8% of female progeny carried *w*^GDe^, indicating a conversion rate of 20% (32/160 p-value: <0.001^***^). This large difference in homing rate between crosses with maternal vs paternal F_0_ *Cas9* suggests that for *bgcn* maternally deposited Cas9 may contribute more to homing than autonomously expressed Cas9. Low expression, but high maternal deposition rate of *bgcn* is also consistent with the difference in the observed *white* phenotype rate (2b). However, the difference in homing events accounts for less than half of the overall difference in *w*^GDe^ inheritance rate between maternally (66%) and paternally (50%) contributed *bgcn Cas9*. This may suggest that another inheritance biasing mechanism is active.

For the *nup50* double heterozygote males with paternal *Cas9* contribution, 12% of their progeny were *w*^GDe^ positive males, indicating 24% (210/869 p-value: <0.001^***^) of the recipient chromosomes were converted by homing. For maternally contributed *Cas9* this was 11% of progeny and 23% (315/1387 p-value: <0.001^***^) of recipient chromosomes. We also performed this analysis on the *nup50* crosses reported in Li et al. (Table S5). Despite a significant increase in the inheritance of the *w*^GDe^ element, there was no evidence of an increased recombination rate: 0% (3/690 p-value: 1.0^ns^) for paternal *Cas9* and 0% (3/688 p-value: 1.0^ns^) for maternal *Cas9* contribution. Instead, there was a significant bias in favour of the *w*^GDe^ linked sex corresponding to the donor chromosome. For paternally contributed *Cas9*, 65% (1306/1996 p-value: <0.001^***^) of progeny were female, >99% of which were *w*^GDe^ positive. For maternally contributed *Cas9* 67% (1371/2059 p-value: <0.001^***^) of progeny were male, >99% of which were *w*^GDe^ positive (Table S16). This sex-bias should not occur through homing. Instead, this seems consistent with a meiotic drive mechanism where some of the non-*w*^GDe^ chromosomes are lost, or conversion of a very large region encompassing both *w*^GDe^ and the sex-determining region (Fig 3a). For the crosses performed for this study, including the *nup50* line, no significant difference in sex, and by extension recipient vs donor chromosome inheritance, was detected (Table S16). For *bgcn* with maternal F_0_ *Cas9*, 59% of all F_2_s were male, but this did not rise to our significance threshold due to the relatively low number of progeny scored for this cross.

## Discussion

In this study, we report the efficiency and mechanisms of three CRISPR-*Cas9* nuclease gene drives targeting the *white* gene, expanding the tool-set for developing genetic control strategies for the public-health relevant *Aedes aegypti* mosquito. In our hands, *sds3, bgcn*, and *nup50* expressed Cas9 each resulted in increased inheritance of the *w*^GDe^ drive element, with the primary mechanism seeming to be homing. In addition, for each promoter, we find evidence of maternal deposition and somatic expression and, unexpectedly, a currently unexplained effect of the *Cas9* carrying grandparent’s sex on the grand-offspring phenotypes (Fig 2b) that we termed Grandparent Enhanced Somatic Phenotype (GESP). In line with Li et al., we find the *white* locus to be a good drive target, allowing for efficient transmission bias and convenient readout of an easily-scored visible recessive phenotype^36^. In addition, the locus allows for effective transgene expression from a sex-linked locus which may be of particular use for future drives and other genetic control approaches. For the *bgcn* drive in males, the recipient chromosome conversion rate was much higher with maternally contributed *Cas9* (19%) compared to paternally contributed *Cas9* (5%). These results suggest that, in at least males, the *bgcn* drive may function primarily through maternally contributed Cas9. Homing through Cas9 deposition in the absence of expressed Cas9 (’shadow drive’) has been reported for other drives^29,30,35^, but to our knowledge not as the primary means of inheritance bias for a drive. We find *nup50* and *sds3*-Cas9 capable of directing transmission bias in females and males, and we did not find that maternal deposition from the *Cas9* carrying grandmother negatively influenced the homing rate observed in males.

For all drives, the almost complete absence of any somatic phenotype in individuals that did not inherit the *w*^GDe^ element (Fig S2) could indicate that, while maternal deposition of the Cas9 occurs, the gRNA^w^ or gRNA^w^:Cas9 complex are either not deposited or rapidly degraded. However, progeny that did not inherit the *w*^GDe^ element instead inherited the (initially) *w*^+^ allele from the double heterozygous parent. For mosaic eye phenotypes to occur in these individuals, up to two functional *w*^+^ alleles may need to be disrupted by deposition instead of one; direct comparison of the rates of somatic mutation between offspring that do and do not inherit the *gRNA*^*w*^ transgene are therefore potentially misleading. Moreover, some non-*w*^GDe^ progeny may have inherited a *white* allele that contained a functional, but cut-resistant, NHEJ mutation (type-1 resistant mutation) which would make bi-allelic disruption impossible. For the − *w*^GDe^ F_2_ progeny, maternal deposition from the F_0_ grandmother could increase their probability of inheriting a mutated *w* allele from their F_1_ parent. As such, GESP does not apply, and only refers to +*w*^GDe^ F_2_ progeny where the sex of the *w*^GDe^ or *Cas9* carrying grandparent influences their propensity to present with a somatic phenotype. While deposition from a F_0_ grandparent may explain a change in the quantity of *w*^GDe^ alleles passed along by the F_1_ drive parent (shadow-homing), it does not seem to explain a change in the phenotype of those F_2_ progeny that inherited a drive element (GESP). Genomic imprinting or transgenerational persistence of the deposited Cas9 mRNA/protein may underlie GESP.

For *nup50* the overall inheritance biasing rate and somatic/embryonic drive activity (*≈*100%) closely match those reported by Li et al.^36^ and underscore its potential utility for systems such as precision-guided SIT^55^. Yet, an important finding of our work is the propensity of this drive to function through two different mechanisms. The selective inheritance or elimination of a chromosome is generally achieved through creating multiple DNA breaks on the target chromosome^10, 56–58^ (e.g. X-shredder) or by disrupting an essential gene^7,8^. Meiotic drive through a single cut in a non-essential gene as found by Li et al., and reported here, is noteworthy. One explanation could be the chromosomal location of the induced double-stranded break. A single cut has been demonstrated to be sufficient for inheritance bias through the loss of a chromosome in yeast when it is targeted to a centromere, while nearby sites were not sufficient^59^. The *white* gene is located relatively near the centromere. However, a centromere effect does not explain the difference in results from this study and that of Li et al., which instead suggests subtle differences in the rearing conditions or background genetics of the mosquito strains may have a significant influence on the underlying mechanisms. Gene drive assessment performed in *D. melanogaster* with different genetic background revealed differences in their activity but changes in the underlying mechanism were not investigated^25^. The *nup50*-*Cas9* and *w*^GDe^ transgenic lines used in this study are the same as described in Li et al., but the crosses to assess homing were made to LVP strains maintained for a long period of time in different insectaries. Mosquito colonies maintained in laboratories can suffer from founder and drift effects, affecting their genetic background and reducing their heterozygosity^60^. Moreover, genetic variability in *A. aegypti* colonies from the same strain but reared in different laboratories has been documented^61^.

We cannot rule out that the sex-bias we report for Li et al.’s *nup50*-*Cas9* is due to copying of a >45 Mbp^48,49^ region comprising both the *w*^GDe^ and the sex-determining region (Fig 3a). While co-conversion of sequences not directly within a drive element has been reported^35,41^, the large distance between the *w*^GDe^ drive and *M* locus leads us to believe this is less plausible. Moreover, similar homing drives have been reported to be sensitive to the alignments of the homology arms^8,20,24^, and several studies have reported partial homing events^22–24^. These partial homing events are seemingly due to sequences in the drive element (such as the gRNA gene) having undesired homology to the recipient chromosome and results in only part of the drive element being copied over. Similarly, a toxin-antidote CRISPR gene drive element was not copied despite targeting a nearby site on the homologous chromosome^8^. These results are inconsistent with a single DNA-break inducing large scale homing beyond the (immediately) adjacent regions of homology.

To our knowledge, for drive designed to function through homing recipient/donor chromosome markers have been used with non-CRISPR nucleases in *D. melanogaster*^38–40^ and *An. gambiae*^41^ and with CRISPR-*Cas9* in *D. melanogaster*^35^, *A. aegypti*^36^ and *Mus musculus*^37^. Collectively, the publications with *D. melanogaster* provide significant evidence against meiotic drive mechanisms contributing significantly to the observed inheritance bias of the ‘homing’ drives examined. Moreover, *nos*-*Cas9* has been reported to cause inheritance bias in many gene drives with a homing drive design^20,22,24–26,29,30,32^ but failed to do so in a meiotic drive design: despite having far higher cleavage activity, a *nos*-*Cas9* X-shredder meiotic drive did not result in sex-bias, while *β* tub85D expressed Cas9 did^58^. There may however be an exception, a study of drives with a homing design noted a reduced transmission of the recipient chromosome for a set of crosses where the split *Cas9* transgene happened to function as a chromosome marker^52^. It should be noted that the drives in this study targeted essential genes, potentially complicating the interpretation of the mechanism of inheritance bias.

In light of our results, re-evaluation of the *A. gambiae* I-*Sce*I gene drive reported by Windbichler et al. may suggest that a meiotic drive effect in homing drive designs is more widespread^41^. Their drive carrying line had a small marker (*Not*I restriction site) located approximately 0.7 kilobases from the I-*Sce*I cut-site on the recipient chromosome, but not on the donor drive chromosome. They reported 86% inheritance of the drive element from heterozygote males. However, drive alleles that included the *Not*I site only accounted for around half the increased drive allele inheritance. The authors attributed this discrepancy to co-conversion, where homing of the drive element also replaced the nearby *Not*I marker. A combined meiotic drive and homing effect would seem to provide an alternative explanation. In the *M. musculus* drive reported by Grunwald et al. the recipient chromosome had a linked coat colour marker that allowed homing events to be precisely tracked^37^. In females, *vasa*-Cre induced CAG-*Cas9* expression and resulted in homing rates of 42% (36/86) and 11% (5/47) depending on the *Cas9* insertion site. In males, no homing was observed with any drive. However, for the *vasa* drives, males passed along the donor drive chromosome to 63% (45/71) and 54% (49/91) of their progeny, potentially indicating a meiotic drive mechanism in that sex. It should be noted that detecting meiotic drive using this method is less sensitive than detecting homing and more progeny would need to be scored to have confidence in this trend. Together, the *A. aegypti*^36^, *An. gambiae*^41^ and *M. musculus*^37^ drives indicate that meiotic drive in drives intended to function through homing may be a widespread occurrence. Distinguishing these mechanisms requires linked markers, and for some organisms, this type of study may be best reserved for drives that after initial tests warrant further development.

Our work further expands the Cas9 expression patterns that have been tested in the context of mosquito gene drives. It is notable that the drives with a homing design reported in the Anopheles mosquitoes *A. gambiae*^16–19^ and *A. stephensi*^13–15^ almost invariably have a dramatically higher conversion rate than those found in *A. aegypti*. It is not clear what underlies this difference. However, the fact that the modest conversion rate for *nup50*-Cas9 males remains stable despite a change in the mechanism may limit the possible explanations. This stability suggests that the factor(s) negatively affecting the conversion rate in *A. aegypti* are not specific to either homing or meiotic drive. Moreover, it also indicates that the difference in conversion rate observed between mosquito species is likely not due to the species favouring one mechanism over the other. Yet, the difference in mechanism between homing and meiotic drive through gamete destruction has important practical implications: First, the loss of gametes through a meiotic-drive mechanism may negatively affect mating competitiveness by lowering the number of viable gametes, though in some cases gametes may be produced in sufficient excess for this not to be significant. The homing mechanism functions through conversion, and should not affect gamete numbers. For the *nup50* meiotic drive reported in Li et al., male *nup50*-*Cas9* fecundity was tested and found to not differ from wildtype^36^. Second, on a ‘per-cut’, basis meiotic drive is moderately less efficient than homing. When meiotic drive removes a non-drive gamete/embryo, it thereby improves the chances of the remaining gametes/embryos. These may, in addition, to drive carrying gametes, include other wildtype and cut-resistant allele carrying gametes that were not destroyed. In contrast, homing converts a non-drive gamete to a drive gamete which does not benefit any of the leftover non-drive gametes making homing more efficient. Third, the linkage between different drive components may change very significantly depending on the mechanisms: for instance, if in a split-drive system the *Cas9* is located near the gRNA element homing would still only increase the number of gRNA alleles, but not the *Cas9* alleles. However, meiotic drive would increase the inheritance of both the gRNA and *Cas9* element. This could theoretically cause a split-drive or daisy-chain drive^62^ to spread more than anticipated. Locating each element on separate chromosomes would prevent this, and our data suggest that this may be a wise precaution to increase the predictability of their invasiveness. Although, if anticipated or identified in early-stage field trails, a meiotic drive induced linkage between elements could also be leveraged, lowering the required release frequencies^63^. Nonetheless, in regards to risk-assessment of rare recombination events, the genomic distance at which two split-drive elements become strongly linked is presumably still much more permissive for a meiotic drive mechanism as opposed to a homing mechanism. Last, in the case of Li et al.’s *white* targeting *A. aegypti* drive, its linkage to the sex-determining locus caused an otherwise neutral replacement drive to act, in males, like a sex-biasing suppression drive. This might be desirable for some applications, but obviously detrimental if the intended application were different. Most of these concerns apply even if the actual mechanism is co-conversion/copy-grafting of a large chromosome segment as opposed to meiotic drive.

## Acknowledgements

S.A.N.V. and L.A. were supported by funding from the Biotechnology and Biological Sciences Research Council (BBSRC) (Grant BB/M011224/1 to S.A.N.V. and Grants BBS/E/I/00007033, BBS/E/I/00007038, and BBS/E/I/00007039 to L.A.). M.A.E.A., E.G., J.A. and L.A. were supported through an award from DARPA’s Safe Genes program to MIT N66001-17-2-4054. O.S.A, N.P.K., and M.L were supported in part by funding from the Defense Advanced Research Project Agency (DARPA) Safe Genes Program Grant HR0011-17-2-0047. The views, opinions and/or findings expressed are those of the author and should not be interpreted as representing the official views or policies of the U.S. Government.

## Author contributions statement

M.A.E.A. and L.A. conceived of the experiments, E.G. conducted the crosses and data collection, S.A.N.V. analysed the results and wrote the manuscript. S.A.N.V. performed the initial analysis that noted the meiotic drive phenomena, and J.A contributed. O.S.A, M.L, and N.P.K. provided transgenic lines. All authors reviewed and provided comments on the manuscript.

## Supplemental information

**Table S1.** Deriving of the *sds3*-*Cas9* line. The number of injected embryo’s (G0), survivors, and number of G1s screened after crossing to LVP.

| Construct | Helper | Embryos injected | G0 survivors (%) | G1s screened | Positive G1s |
| --- | --- | --- | --- | --- | --- |
| *sds3-Cas9* | Pub-HYPpbac | 990 | 161 (16%) | 4664 | 464 (8/8 pools) |

**Table S2.** F_2_ progeny of *sds3*-*Cas9* and *w*^GDe^ double heterozygotes crossed to LVP wildtype. Mosaic eyes (ME), white eyes (WE). Drive F_1_ phenotypes are shown in Fig S1.

| *sds3* | F1s crossed (♂ x ♀) | Drive F1 phenotype | +w<sup>GDe</sup>;+*Cas9* |  |  |  | +w<sup>GDe</sup>;– |  |  |  | –/+*Cas9* |  |  |  | –/– |  |  |  |
| --- | --- | --- | --- | --- | --- | --- | --- | --- | --- | --- | --- | --- | --- | --- | --- | --- | --- | --- |
| Cross |  |  | ♀ |  | ♂ |  | ♀ |  | ♂ |  | ♀ |  | ♂ |  | ♀ |  | ♂ |  |
|  |  |  | ME/WE | WT | ME/WE | WT | ME/WE | WT | ME/WE | WT | ME/WE | WT | ME/WE | WT | ME/WE | WT | ME/WE | WT |
| F1: ♂w<sup>GDe</sup>;sds3-*Cas9* x ♀LVP<br>F0: ♂*Cas9* x ♀w<sup>GDe</sup> | 10 x 50 | Mosaic eyes | 91 | 13 | 20 | 5 | 0 | 130 | 1 | 15 | 0 | 0 | 0 | 76 | 0 | 0 | 0 | 99 |
| F1: ♂LVP x ♀w<sup>GDe</sup>;sds3-*Cas9*<br>F0: ♂*Cas9* x ♀w<sup>GDe</sup> | 10 x 25 | Mosaic eyes | 38 | 2 | 34 | 9 | 36 | 1 | 39 | 0 | 0 | 22 | 0 | 23 | 0 | 33 | 0 | 38 |
| F1: ♂w<sup>GDe</sup>;sds3-*Cas9* x ♀LVP<br>F0: ♂w<sup>GDe</sup> x ♀*Cas9* | 10 x 50 | Mosaic eyes | 4 | 1 | 57 | 0 | 0 | 14 | 0 | 55 | 1 | 40 | 0 | 0 | 0 | 42 | 0 | 0 |
| F1: ♂LVP x ♀w<sup>GDe</sup>;sds3-*Cas9*<br>F0: ♂w<sup>GDe</sup> x ♀*Cas9* | 10 x 21 | Mosaic eyes | 33 | 1 | 20 | 3 | 28 | 0 | 33 | 0 | 0 | 12 | 0 | 16 | 0 | 12 | 1 | 17 |

**Table S3.**
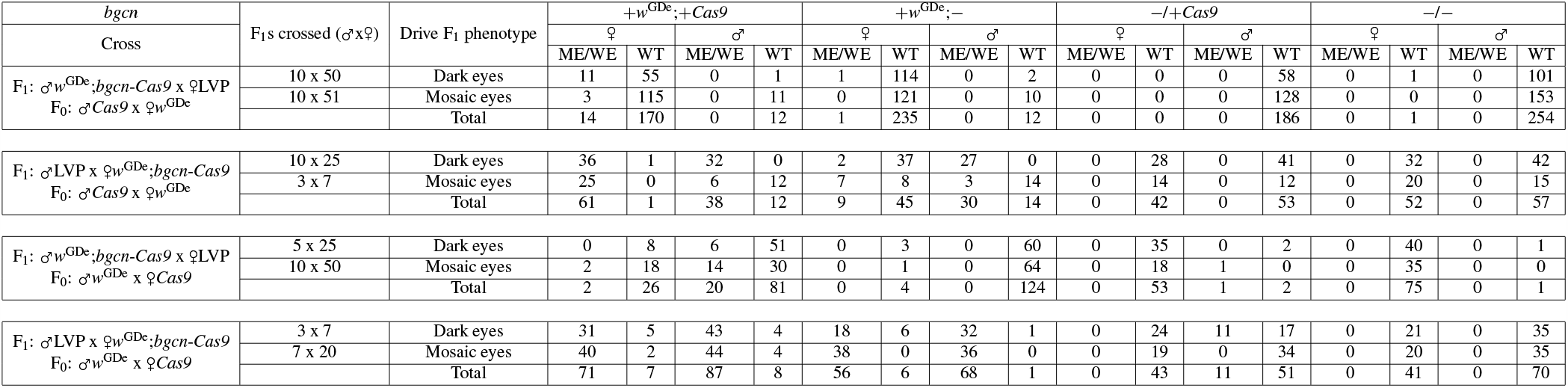
F_2_ progeny of *bgcn*-*Cas9* and *w*^GDe^ double heterozygotes crossed to LVP wildtype. Mosaic eyes (ME), white eyes (WE)

| *bgn* | F<sub>1</sub>s crossed (♂x♀) | Drive F<sub>1</sub> phenotype | +w<sup>GDe</sup>;+Cas9 |  |  |  | +w<sup>GDe</sup>;– |  |  |  | –/+Cas9 |  |  |  | –/– |  |  |  |
| --- | --- | --- | --- | --- | --- | --- | --- | --- | --- | --- | --- | --- | --- | --- | --- | --- | --- | --- |
| Cross |  |  | ♀ |  | ♂ |  | ♀ |  | ♂ |  | ♀ |  | ♂ |  | ♀ |  | ♂ |  |
|  |  |  | ME/WE | WT | ME/WE | WT | ME/WE | WT | ME/WE | WT | ME/WE | WT | ME/WE | WT | ME/WE | WT | ME/WE | WT |
| F<sub>1</sub>: ♂w<sup>GDe</sup>;bgn-Cas9 x ♀LVP<br>F<sub>0</sub>: ♂Cas9 x ♀w<sup>GDe</sup> | 10 x 50 | Dark eyes | 11 | 55 | 0 | 1 | 1 | 114 | 0 | 2 | 0 | 0 | 0 | 58 | 0 | 1 | 0 | 101 |
|  | 10 x 51 | Mosaic eyes | 3 | 115 | 0 | 11 | 0 | 121 | 0 | 10 | 0 | 0 | 0 | 128 | 0 | 0 | 0 | 153 |
|  |  | Total | 14 | 170 | 0 | 12 | 1 | 235 | 0 | 12 | 0 | 0 | 0 | 186 | 0 | 1 | 0 | 254 |
| F<sub>1</sub>: ♂LVP x ♀w<sup>GDe</sup>;bgn-Cas9<br>F<sub>0</sub>: ♂Cas9 x ♀w<sup>GDe</sup> | 10 x 25 | Dark eyes | 36 | 1 | 32 | 0 | 2 | 37 | 27 | 0 | 0 | 28 | 0 | 41 | 0 | 32 | 0 | 42 |
|  | 3 x 7 | Mosaic eyes | 25 | 0 | 6 | 12 | 7 | 8 | 3 | 14 | 0 | 14 | 0 | 12 | 0 | 20 | 0 | 15 |
|  |  | Total | 61 | 1 | 38 | 12 | 9 | 45 | 30 | 14 | 0 | 42 | 0 | 53 | 0 | 52 | 0 | 57 |
| F<sub>1</sub>: ♂w<sup>GDe</sup>;bgn-Cas9 x ♀LVP<br>F<sub>0</sub>: ♂w<sup>GDe</sup> x ♀Cas9 | 5 x 25 | Dark eyes | 0 | 8 | 6 | 51 | 0 | 3 | 0 | 60 | 0 | 35 | 0 | 2 | 0 | 40 | 0 | 1 |
|  | 10 x 50 | Mosaic eyes | 2 | 18 | 14 | 30 | 0 | 1 | 0 | 64 | 0 | 18 | 1 | 0 | 0 | 35 | 0 | 0 |
|  |  | Total | 2 | 26 | 20 | 81 | 0 | 4 | 0 | 124 | 0 | 53 | 1 | 2 | 0 | 75 | 0 | 1 |
| F<sub>1</sub>: ♂LVP x ♀w<sup>GDe</sup>;bgn-Cas9<br>F<sub>0</sub>: ♂w<sup>GDe</sup> x ♀Cas9 | 3 x 7 | Dark eyes | 31 | 5 | 43 | 4 | 18 | 6 | 32 | 1 | 0 | 24 | 11 | 17 | 0 | 21 | 0 | 35 |
|  | 7 x 20 | Mosaic eyes | 40 | 2 | 44 | 4 | 38 | 0 | 36 | 0 | 0 | 19 | 0 | 34 | 0 | 20 | 0 | 35 |
|  |  | Total | 71 | 7 | 87 | 8 | 56 | 6 | 68 | 1 | 0 | 43 | 11 | 51 | 0 | 41 | 0 | 70 |

**Table S4.**
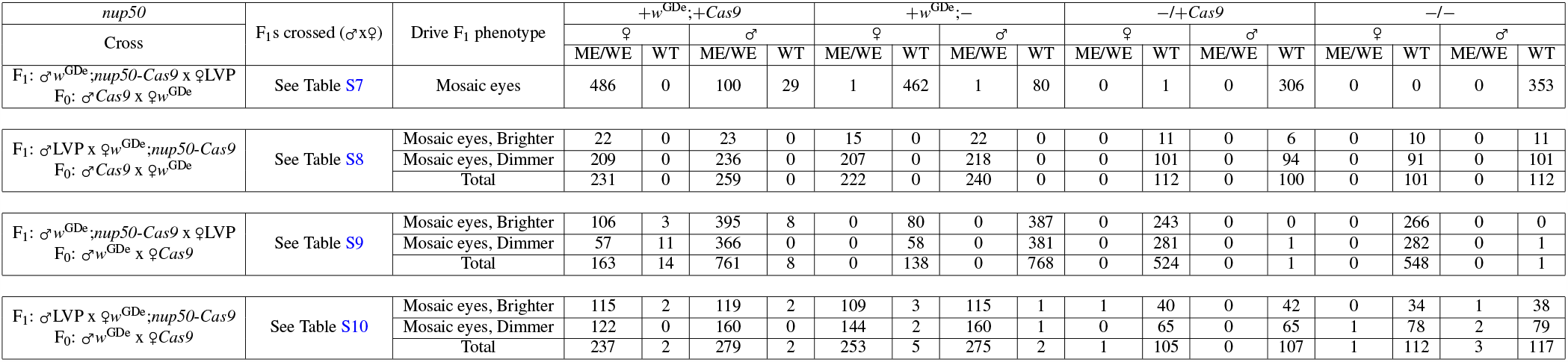
F_2_ progeny of *nup50*-*Cas9* and *w*^GDe^ double heterozygotes crossed to LVP wildtype. Mosaic eyes (ME), white eyes (WE). Drive F_1_ phenotypes are shown in Fig S1.

**Table S5.**
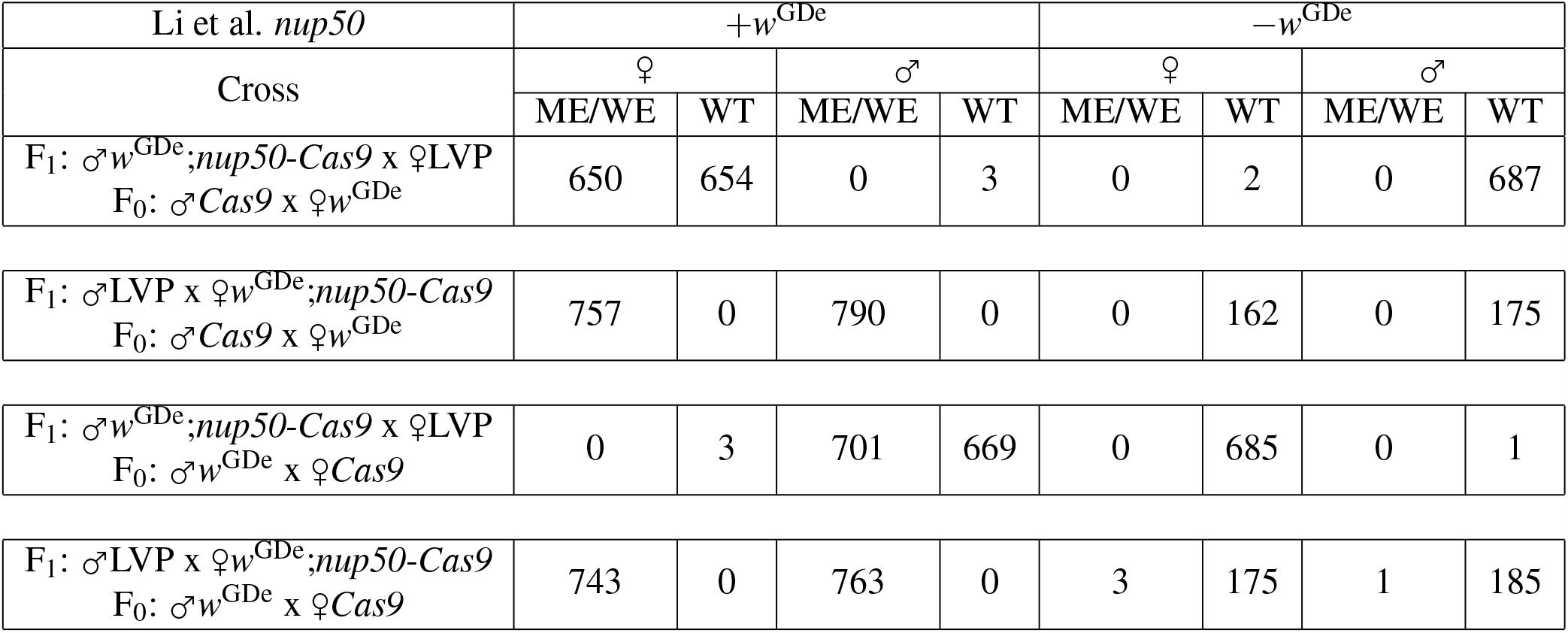
F_2_ progeny of *nup50*-*Cas9* and *w*^GDe^ double heterozygotes crossed to wildtype taken from Li et al. Mosaic eyes (ME), white eyes (WE). Data are, in row order, from Li et al. supplemental files 4f, 4e, 4d, and 4c.

| Li et al. *nup50* | $+w^{GDe}$ | | | | $-w^{GDe}$ | | | |
| --- | --- | --- | --- | --- | --- | --- | --- | --- |
| Cross | ♀ |  | ♂ |  | ♀ |  | ♂ |  |
|  | ME/WE | WT | ME/WE | WT | ME/WE | WT | ME/WE | WT |
| F<sub>1</sub>: ♂ $w^{GDe};nup50-Cas9$ x ♀ LVP<br>F<sub>0</sub>: ♂ *Cas9* x ♀ $w^{GDe}$ | 650 | 654 | 0 | 3 | 0 | 2 | 0 | 687 |
| F<sub>1</sub>: ♂ LVP x ♀ $w^{GDe};nup50-Cas9$<br>F<sub>0</sub>: ♂ *Cas9* x ♀ $w^{GDe}$ | 757 | 0 | 790 | 0 | 0 | 162 | 0 | 175 |
| F<sub>1</sub>: ♂ $w^{GDe};nup50-Cas9$ x ♀ LVP<br>F<sub>0</sub>: ♂ $w^{GDe}$ x ♀ *Cas9* | 0 | 3 | 701 | 669 | 0 | 685 | 0 | 1 |
| F<sub>1</sub>: ♂ LVP x ♀ $w^{GDe};nup50-Cas9$<br>F<sub>0</sub>: ♂ $w^{GDe}$ x ♀ *Cas9* | 743 | 0 | 763 | 0 | 3 | 175 | 1 | 185 |

**Table S6.** F_2_ progeny of *w*^GDe^ heterozygotes (no *Cas9*) crossed to LVP wildtype. Mosaic eyes (ME), white eyes (WE)

| *Cas9* | Drive F<sub>1</sub> phenotype | *+w<sup>GDe</sup>; +Cas9* |  |  |  | *+w<sup>GDe</sup>; -* |  |  |  | *-/+Cas9* |  |  |  | *-/-* |  |  |  |
| --- | --- | --- | --- | --- | --- | --- | --- | --- | --- | --- | --- | --- | --- | --- | --- | --- | --- |
| Cross |  | ♀ |  | ♂ |  | ♀ |  | ♂ |  | ♀ |  | ♂ |  | ♀ |  | ♂ |  |
|  |  | ME/WE | WT | ME/WE | WT | ME/WE | WT | ME/WE | WT | ME/WE | WT | ME/WE | WT | ME/WE | WT | ME/WE | WT |
| F<sub>1</sub>: ♂ *w<sup>GDe</sup>* x ♀ LVP<br>F<sub>0</sub>: ♂ *w<sup>GDe</sup>* x ♀ LVP | Dark eyes | 0 | 0 | 0 | 0 | 0 | 11 | 0 | 609 | 0 | 0 | 0 | 0 | 0 | 581 | 0 | 2 |
| F<sub>1</sub>: ♂ LVP x ♀ *w<sup>GDe</sup>*<br>F<sub>0</sub>: ♂ *w<sup>GDe</sup>* x ♀ LVP | Dark eyes | 0 | 0 | 0 | 0 | 0 | 145 | 0 | 163 | 0 | 0 | 0 | 0 | 0 | 147 | 0 | 150 |

**Table S7.**
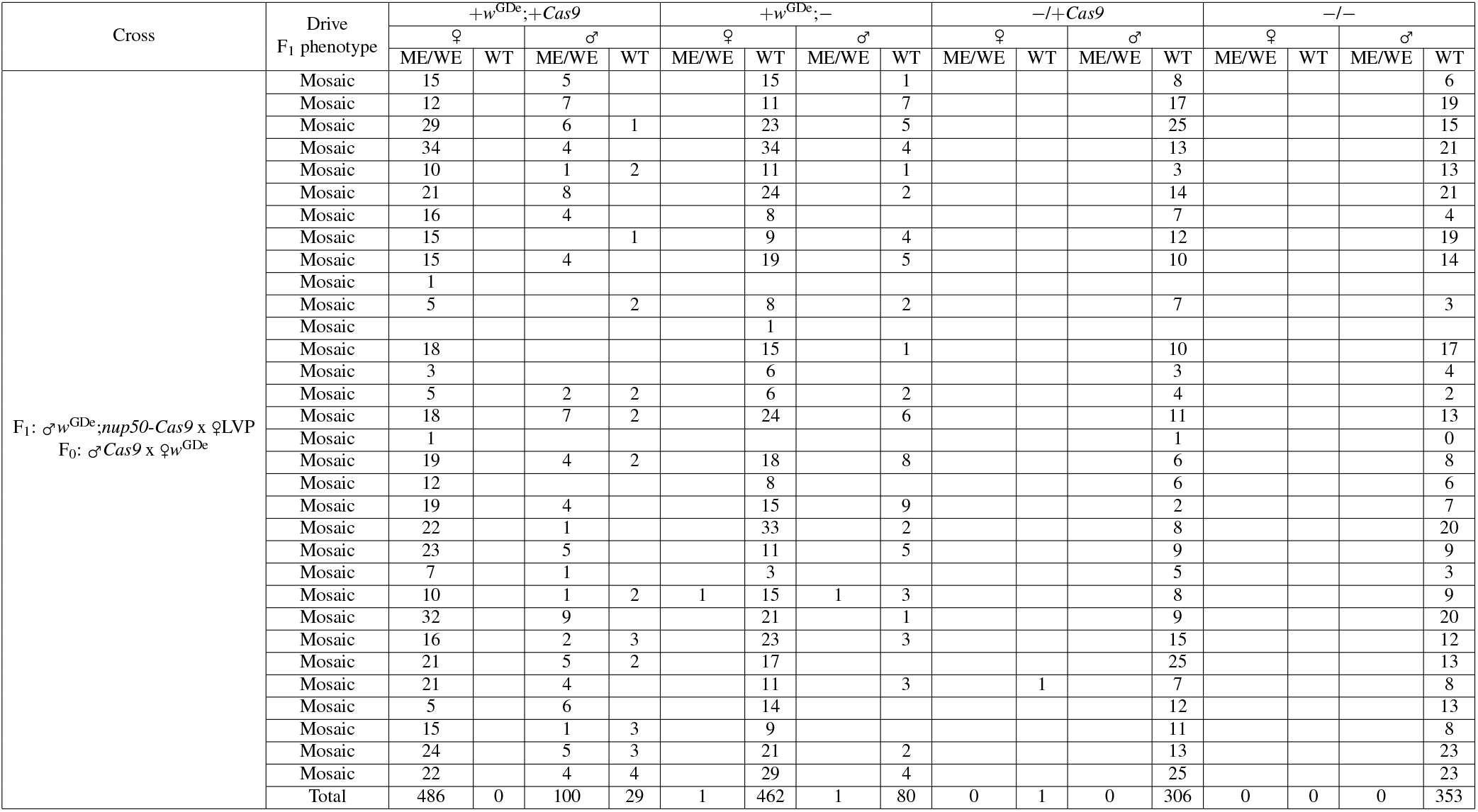
F_2_ progeny of individual male *nup50*-*Cas9* and *w*^GDe^ double heterozygotes crossed to LVP wildtype. *nup50*-*Cas9* from paternal F_0_. Mosaic eyes (ME), white eyes (WE). Drive F_1_ phenotypes are shown in Fig S1.

**Table S8.** F_2_ progeny of individual female *nup50*-*Cas9* and *w*^GDe^ double heterozygotes crossed to LVP wildtype. *nup50*-*Cas9* from paternal F_0_. Mosaic eyes (ME), white eyes (WE). Drive F_1_ phenotypes are shown in Fig S1.

| Cross | Drive<br>F<sub>1</sub> phenotype | +w<sup>GDe</sup>;+Cas9 |  |  |  | +w<sup>GDe</sup>;- |  |  |  | -/+Cas9 |  |  |  | -/- |  |  |  |
| --- | --- | --- | --- | --- | --- | --- | --- | --- | --- | --- | --- | --- | --- | --- | --- | --- | --- |
|  |  | ♀ |  | ♂ |  | ♀ |  | ♂ |  | ♀ |  | ♂ |  | ♀ |  | ♂ |  |
|  |  | ME/WE | WT | ME/WE | WT | ME/WE | WT | ME/WE | WT | ME/WE | WT | ME/WE | WT | ME/WE | WT | ME/WE | WT |
| F<sub>1</sub>: ♂ LVP x ♀ w<sup>GDe</sup>;nup50-Cas9<br>F<sub>0</sub>: ♂ Cas9 x ♀ w<sup>GDe</sup> | Mosaic, Dimmer | 6 |  | 5 |  | 8 |  | 2 |  |  | 3 |  | 4 |  | 2 |  | 2 |
|  | Mosaic, Dimmer | 10 |  | 9 |  | 10 |  | 7 |  |  | 1 |  | 3 |  | 3 |  | 4 |
|  | Mosaic, Dimmer | 7 |  | 6 |  | 7 |  | 6 |  |  | 4 |  |  |  | 4 |  | 5 |
|  | Mosaic, Dimmer | 20 |  | 12 |  | 0 |  | 11 |  |  |  |  | 5 |  | 3 |  | 4 |
|  | Mosaic, Dimmer | 7 |  | 12 |  | 8 |  | 15 |  |  | 5 |  | 4 |  | 5 |  | 5 |
|  | Mosaic, Dimmer | 5 |  | 15 |  | 9 |  | 20 |  |  | 5 |  |  |  | 9 |  | 4 |
|  | Mosaic, Dimmer | 12 |  | 14 |  | 11 |  | 10 |  |  | 8 |  | 5 |  | 4 |  | 8 |
|  | Mosaic, Dimmer | 3 |  |  |  | 1 |  |  |  |  |  |  |  |  | 1 |  |  |
|  | Mosaic, Dimmer | 3 |  | 4 |  | 4 |  | 6 |  |  | 8 |  | 3 |  | 4 |  | 3 |
|  | Mosaic, Dimmer | 7 |  | 6 |  | 2 |  | 6 |  |  | 4 |  | 4 |  | 6 |  | 5 |
|  | Mosaic, Dimmer | 9 |  | 6 |  | 13 |  | 7 |  |  | 2 |  | 3 |  | 4 |  | 8 |
|  | Mosaic, Dimmer |  |  |  |  | 2 |  |  |  |  |  |  |  |  |  |  |  |
|  | Mosaic, Dimmer | 5 |  | 9 |  | 17 |  | 13 |  |  | 10 |  | 5 |  | 3 |  | 4 |
|  | Mosaic, Dimmer | 17 |  | 16 |  | 9 |  | 14 |  |  | 5 |  | 4 |  | 3 |  | 4 |
|  | Mosaic, Dimmer | 7 |  | 17 |  | 5 |  | 12 |  |  | 3 |  | 5 |  | 5 |  | 5 |
|  | Mosaic, Dimmer | 4 |  | 8 |  | 13 |  | 2 |  |  | 2 |  | 7 |  | 4 |  | 7 |
|  | Mosaic, Dimmer | 3 |  | 11 |  | 3 |  | 2 |  |  | 6 |  | 4 |  | 1 |  | 2 |
|  | Mosaic, Dimmer | 13 |  | 5 |  | 10 |  | 5 |  |  | 5 |  | 2 |  | 6 |  | 3 |
|  | Mosaic, Dimmer | 19 |  | 16 |  | 7 |  | 8 |  |  | 6 |  | 6 |  | 3 |  | 4 |
|  | Mosaic, Dimmer | 4 |  | 4 |  | 8 |  | 6 |  |  | 6 |  | 6 |  | 5 |  | 5 |
|  | Mosaic, Dimmer | 6 |  | 5 |  | 3 |  | 5 |  |  | 2 |  | 1 |  | 6 |  | 2 |
|  | Mosaic, Dimmer | 12 |  | 15 |  | 18 |  | 14 |  |  | 12 |  | 7 |  | 3 |  | 2 |
|  | Mosaic, Dimmer | 9 |  | 15 |  | 16 |  | 18 |  |  | 2 |  | 7 |  | 2 |  | 3 |
|  | Mosaic, Dimmer | 9 |  | 10 |  | 8 |  | 11 |  |  |  |  | 2 |  |  |  | 4 |
|  | Mosaic, Dimmer | 12 |  | 16 |  | 15 |  | 18 |  |  | 2 |  | 7 |  | 5 |  | 8 |
|  | Mosaic, Brighter | 5 |  | 7 |  | 4 |  | 6 |  |  | 1 |  | 3 |  | 3 |  | 4 |
|  | Mosaic, Brighter | 6 |  | 5 |  | 4 |  | 1 |  |  | 2 |  |  |  | 3 |  | 3 |
|  | Mosaic, Brighter | 8 |  | 11 |  | 5 |  | 14 |  |  | 6 |  | 2 |  | 3 |  | 3 |
|  | Mosaic, Brighter | 2 |  |  |  |  |  |  |  |  |  |  |  |  |  |  |  |
|  | Mosaic, Brighter | 1 |  |  |  | 2 |  | 1 |  |  | 2 |  | 1 |  | 1 |  | 1 |
|  | Sum | 231 | 0 | 259 | 0 | 222 | 0 | 240 | 0 | 0 | 112 | 0 | 100 | 0 | 101 | 0 | 112 |

**Table S9.**
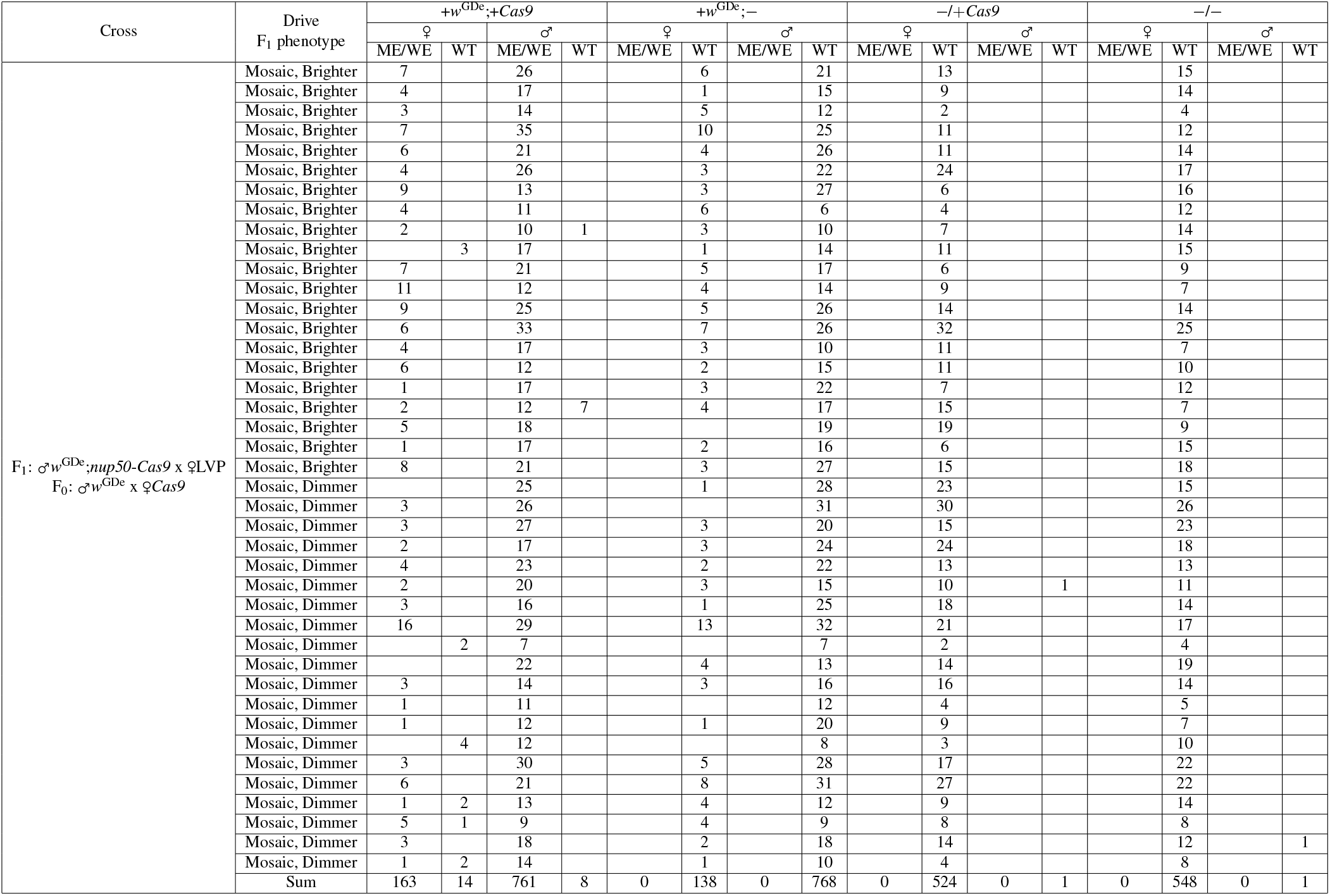
F_2_ progeny of individual male *nup50*-*Cas9* and *w*^GDe^ double heterozygotes crossed to LVP wildtype. *nup50*-*Cas9* from maternal F_0_. Mosaic eyes (ME), white eyes (WE). Drive F_1_ phenotypes are shown in Fig S1.

**Table S10.**
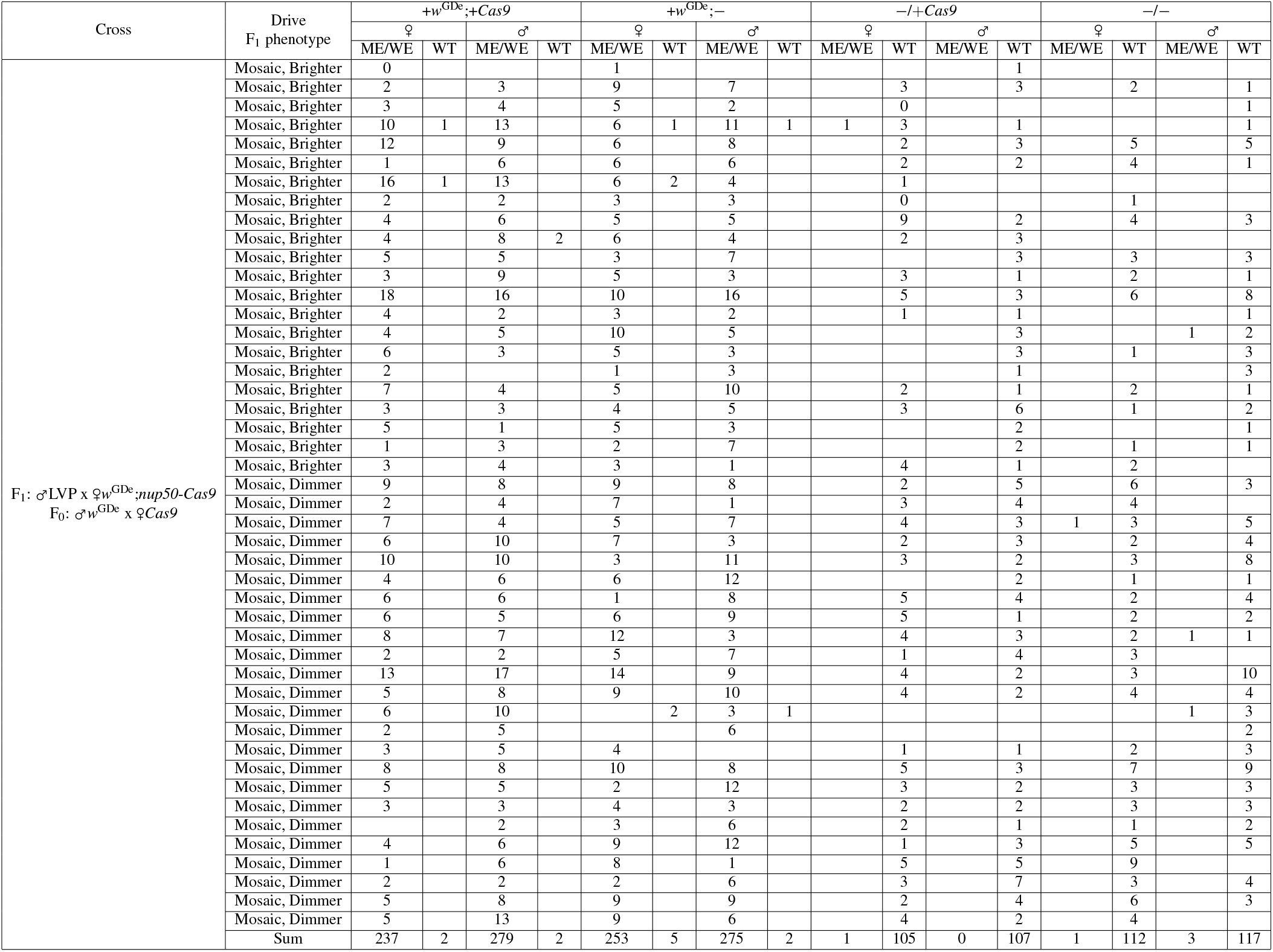
F_2_ progeny of individual female *nup50*-*Cas9* and *w*^GDe^ double heterozygotes crossed to LVP wildtype. *nup50*-*Cas9* from maternal F_0_. Mosaic eyes (ME), white eyes (WE). Drive F_1_ phenotypes are shown in Fig S1.

**Table S11.** Overall *w*^GDe^ inheritance bias. in each case, a Fisher’s exact test is performed using the -*Cas9* condition with the matching drive F_1_ sex. Significance thresholds: * for <=0.05, ** for <=0.01, *** for <=0.001.

| Cas9 | Cas9 F<sub>0</sub> | Drive F<sub>1</sub> | +w<sup>GDe</sup> F<sub>2</sub> | Total F<sub>2</sub> | % | p-value | Threshold | Odds ratio |
| --- | --- | --- | --- | --- | --- | --- | --- | --- |
| -Cas9 | N/A | ♂ | 620 | 1203 | 51.5% | 1.00E+00 | ns | 1 |
| -Cas9 | N/A | ♀ | 308 | 605 | 50.9% | 1.00E+00 | ns | 1 |
| *sds3* | ♂ | ♂ | 275 | 450 | 61.1% | 6.58E-02 | ns | 1.2 |
| *sds3* | ♀ | ♂ | 131 | 214 | 61.2% | 1.57E-01 | ns | 1.2 |
| *sds3* | ♂ | ♀ | 159 | 275 | 57.8% | 2.98E-01 | ns | 1.1 |
| *sds3* | ♀ | ♀ | 118 | 176 | 67.0% | 4.95E-02 | * | 1.3 |
| *bgn* | ♂ | ♂ | 444 | 885 | 50.2% | 7.32E-01 | ns | 1.0 |
| *bgn* | ♀ | ♂ | 257 | 389 | 66.1% | 9.75E-03 | ** | 1.3 |
| *bgn* | ♂ | ♀ | 210 | 414 | 50.7% | 1.00E+00 | ns | 1.0 |
| *bgn* | ♀ | ♀ | 304 | 520 | 58.5% | 1.75E-01 | ns | 1.1 |
| *nup50* | ♂ | ♂ | 1159 | 1819 | 63.7% | 6.32E-04 | *** | 1.2 |
| *nup50* | ♀ | ♂ | 1852 | 2926 | 63.3% | 3.76E-04 | *** | 1.2 |
| *nup50* | ♂ | ♀ | 952 | 1377 | 69.1% | 1.67E-04 | *** | 1.4 |
| *nup50* | ♀ | ♀ | 1055 | 1501 | 70.3% | 6.61E-05 | *** | 1.4 |

**Table S12.**
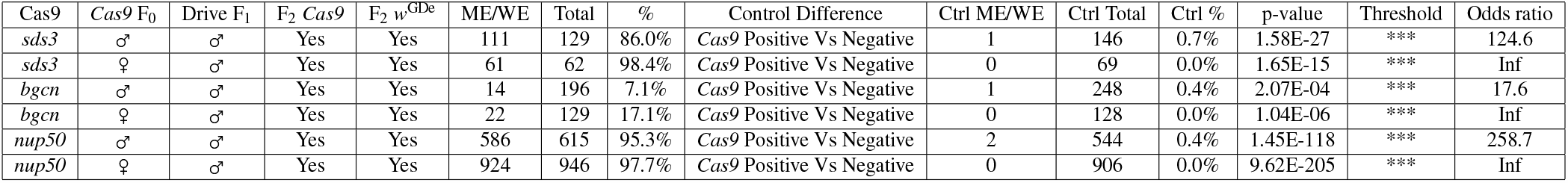
Fisher’s exact test for somatic expression. Mosaic eyes (ME), white eyes (WE). Significance thresholds: * for <=0.05, ** for <=0.01, *** for <=0.001.

| Cas9 | Cas9 F<sub>0</sub> | Drive F<sub>1</sub> | F<sub>2</sub> Cas9 | F<sub>2</sub> w<sup>GDe</sup> | ME/WE | Total | % | Control Difference | Ctrl ME/WE | Ctrl Total | Ctrl % | p-value | Threshold | Odds ratio |
| --- | --- | --- | --- | --- | --- | --- | --- | --- | --- | --- | --- | --- | --- | --- |
| *sds3* | ♂ | ♂ | Yes | Yes | 111 | 129 | 86.0% | Cas9 Positive Vs Negative | 1 | 146 | 0.7% | 1.58E-27 | *** | 124.6 |
| *sds3* | ♀ | ♂ | Yes | Yes | 61 | 62 | 98.4% | Cas9 Positive Vs Negative | 0 | 69 | 0.0% | 1.65E-15 | *** | Inf |
| *bgn* | ♂ | ♂ | Yes | Yes | 14 | 196 | 7.1% | Cas9 Positive Vs Negative | 1 | 248 | 0.4% | 2.07E-04 | *** | 17.6 |
| *bgn* | ♀ | ♂ | Yes | Yes | 22 | 129 | 17.1% | Cas9 Positive Vs Negative | 0 | 128 | 0.0% | 1.04E-06 | *** | Inf |
| *nup50* | ♂ | ♂ | Yes | Yes | 586 | 615 | 95.3% | Cas9 Positive Vs Negative | 2 | 544 | 0.4% | 1.45E-118 | *** | 258.7 |
| *nup50* | ♀ | ♂ | Yes | Yes | 924 | 946 | 97.7% | Cas9 Positive Vs Negative | 0 | 906 | 0.0% | 9.62E-205 | *** | Inf |

**Table S13.** Fisher’s exact test for maternal deposition. Mosaic eyes (ME), white eyes (WE). Significance thresholds: * for <=0.05, ** for <=0.01, *** for <=0.001.

| Cas9 | Cas9 F<sub>0</sub> | Drive F<sub>1</sub> | F<sub>2</sub> Cas9 | F<sub>2</sub> w<sup>GDe</sup> | ME/WE | Total | % | Control Difference | Ctrl ME/WE | Ctrl Total | Ctrl % | p-value | Threshold | Odds ratio |
| --- | --- | --- | --- | --- | --- | --- | --- | --- | --- | --- | --- | --- | --- | --- |
| *sds3* | ♂ | ♀ | No | Yes | 75 | 76 | 98.7% | Drive F<sub>1</sub> ♀ vs ♂ | 1 | 146 | 0.7% | 1.80E-26 | *** | 141.9 |
| *sds3* | ♀ | ♀ | No | Yes | 61 | 61 | 100.0% | Drive F<sub>1</sub> ♀ vs ♂ | 0 | 69 | 0.0% | 7.36E-16 | *** | Inf |
| *bgn* | ♂ | ♀ | No | Yes | 39 | 98 | 39.8% | Drive F<sub>1</sub> ♀ vs ♂ | 1 | 248 | 0.4% | 1.49E-18 | *** | 97.9 |
| *bgn* | ♀ | ♀ | No | Yes | 124 | 131 | 94.7% | Drive F<sub>1</sub> ♀ vs ♂ | 0 | 128 | 0.0% | 1.61E-28 | *** | Inf |
| *nup50* | ♂ | ♀ | No | Yes | 462 | 462 | 100.0% | Drive F<sub>1</sub> ♀ vs ♂ | 2 | 544 | 0.4% | 2.29E-115 | *** | 271.3 |
| *nup50* | ♀ | ♀ | No | Yes | 528 | 535 | 98.7% | Drive F<sub>1</sub> ♀ vs ♂ | 0 | 906 | 0.0% | 8.43E-178 | *** | Inf |

**Figure S1.**
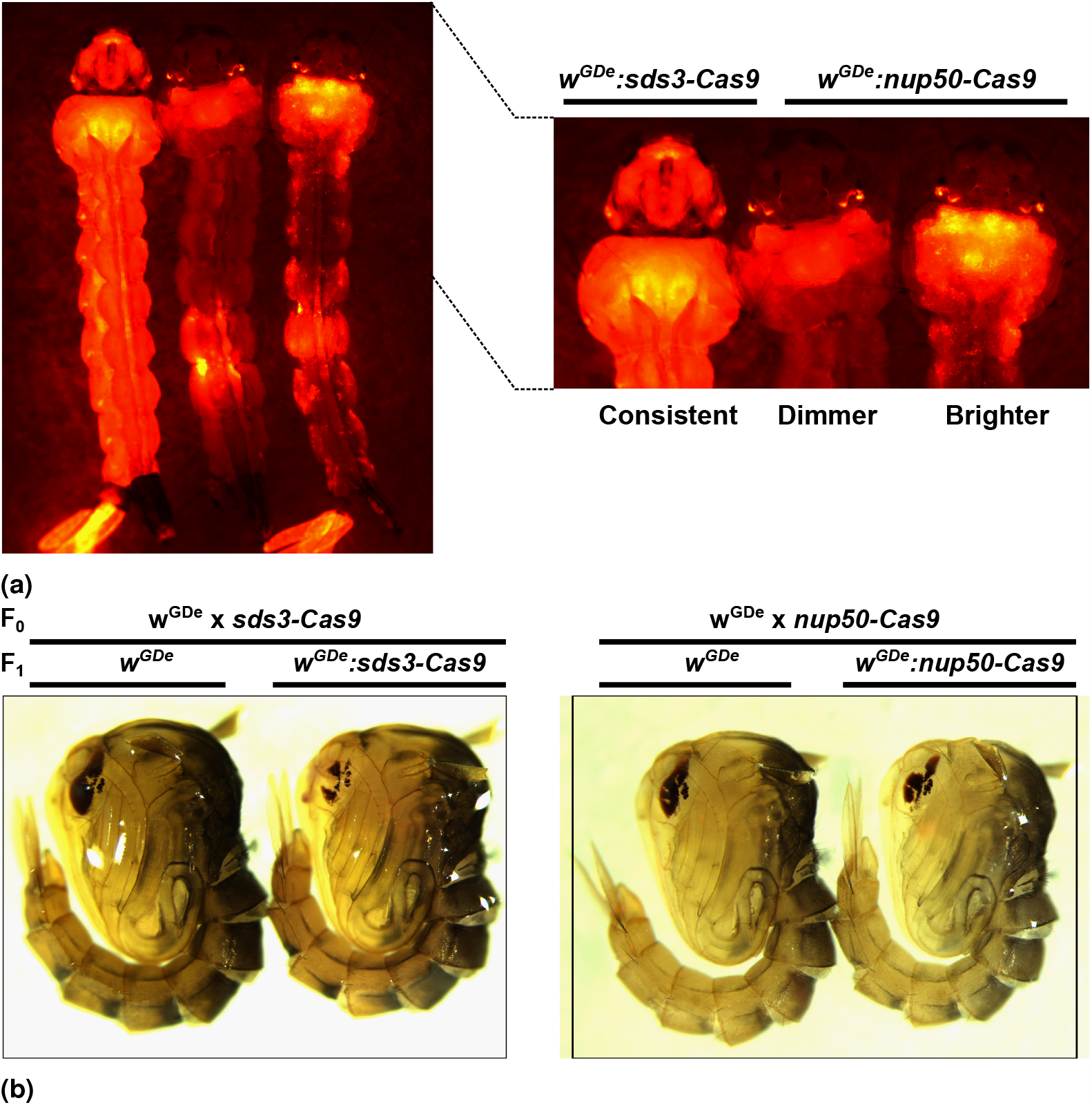
Fluorescent marker phenotype and *white* gene eye phenotype noted among the F_1_s. a ‘Dimmer’ and ‘Brighter’ OpIE2-DsRED phenotype noted in the *nup50*-*Cas9* line. The *sds3* and *bgcn*-*Cas9* used PUb-mCherry-SV40 and had a consistent phenotype. b Examples of the F_1_ dark-eyed (WT) and mosaic eyed phenotype.

**Figure S2.**
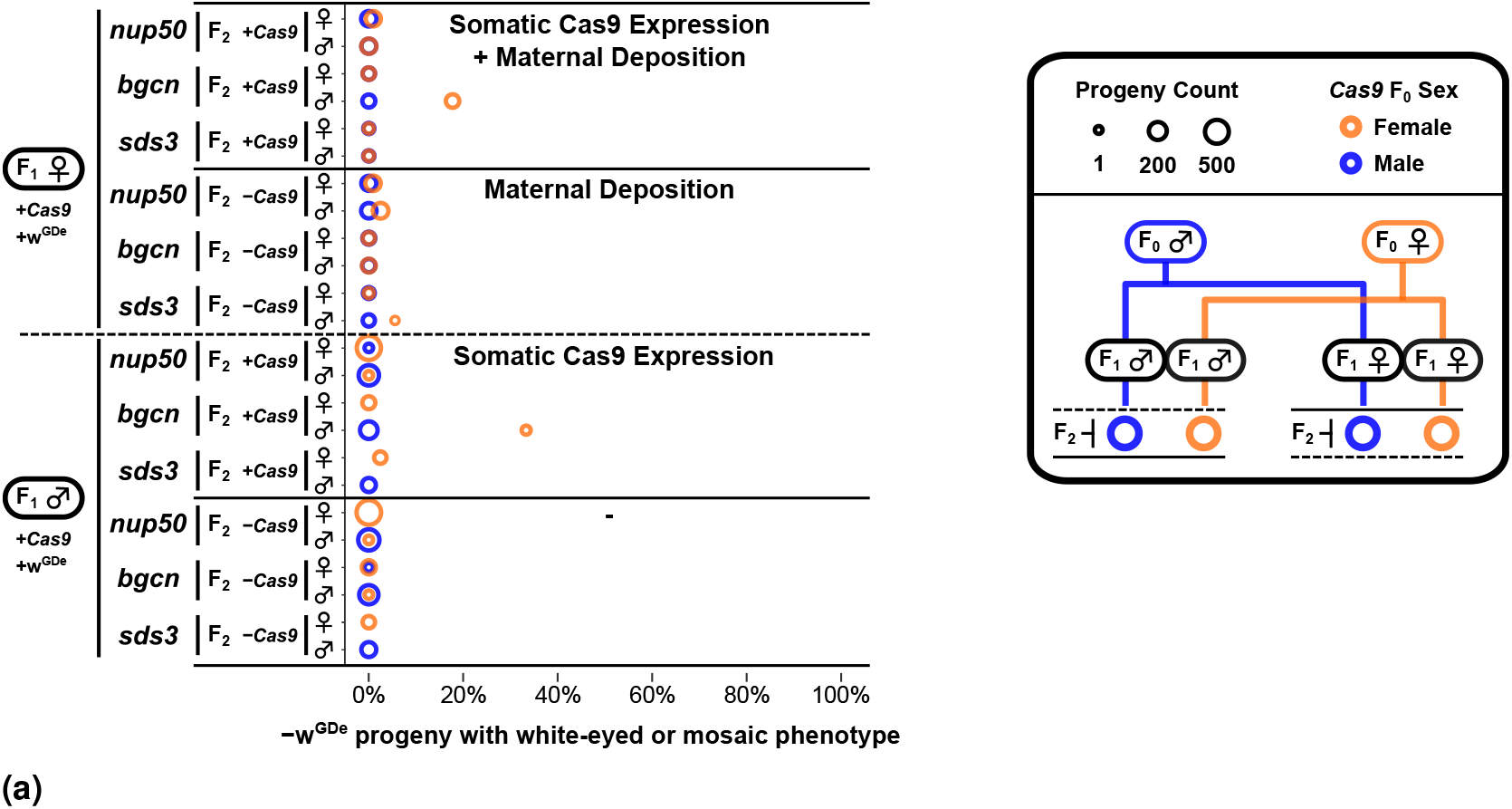
Somatic eye phenotype in the − *w*^GDe^ progeny of double heterozygote split-drive carriers. The percentage of − *w*^GDe^ progeny that display a mosaic or total loss of eye pigment phenotype. F_2_ progeny are segregated by the drive carrying F_1_’s sex (♂,♀), the F_2_’s *Cas9* transgene inheritance (− *Cas9*, +*Cas9*), the *Cas9* regulatory sequences (*sds3, bgcn, nup50*), and the F_2_’s sex (♂,♀). The circle size indicates the number of progeny that make up that group, and circle colour indicates if the *Cas9* carrying F_0_ grandparent was male (Blue) or female (Orange). The set of progeny that came from drive F_1_ carrying females are indicated with ‘Maternal Deposition’. The set of progeny that inherited a *Cas9* element are indicated with ‘Somatic Cas9 Expression’. Due to the sex-linkage of the *w*^GDe^ element from F_1_ drive males the number of F_2_ progeny are unequally distributed among the groups for each sex. This sex-bias is more pronounced for the − *w*^GDe^ progeny than for the +*w*^GDe^ progeny due to the the effects of homing. Groups with <1 progeny are not shown. Within matched crosses (each row), differences in the white phenotype rate corresponding to the *Cas9* carrying F_0_’s sex cannot be solely attributed to a grandparent enhanced somatic phenotype for − *w*^GDe^ progeny. White phenotype rates for +*w*^GDe^ progeny are shown in Fig 2b. Note that inheritance of the *w*^GDe^ element also prevents the potential inheritance from the drive parent of an undamaged wildtype *white* gene or a *white* gene mutation that retains its function (r1) in pigment production. As such, any difference in white phenotype rates between +*w*^GDe^ and − *w*^GDe^ progeny cannot be solely attributed to gRNA expression from *w*^GDe^ in the F_2_ progeny.

**Table S14.**
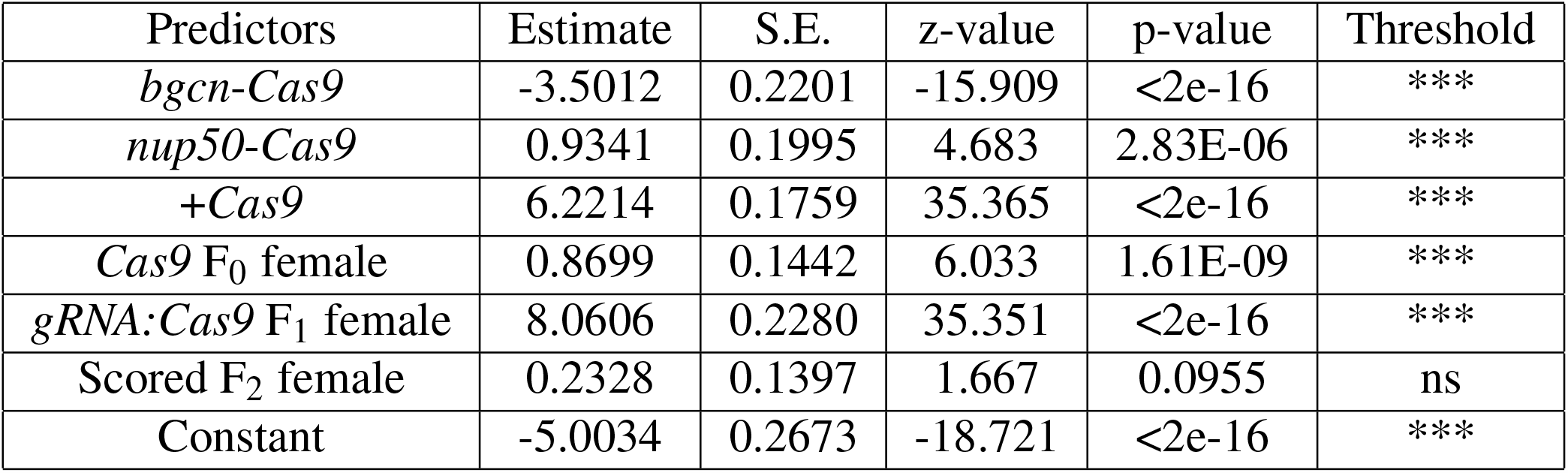
The results of the binomial generalised linear model applied to mosaic eyed (ME) or white-eyed (WE) phenotype fraction among all +*w*^GDe^ F_2_ progeny in Table S2-S4. No interaction terms were specified. The *sds3*-*Cas9*, -*Cas9, Cas9* F_0_ male, *gRNA:Cas9* F_1_ male, F_2_ male cross serves as the reference result. Significance thresholds: * for <=0.05, ** for <=0.01, *** for <=0.001.

**Table S15.**
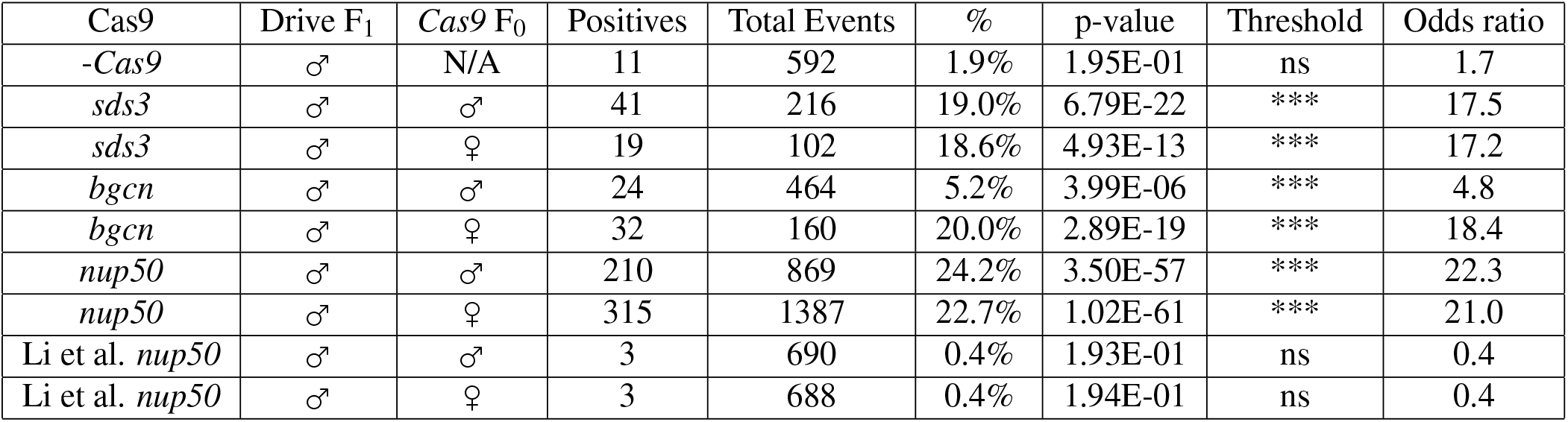
Homing significance test based on the recombination of the +*w*^GDe^ allele and sex-determining region. The total events are all female F_2_s for the ♀ *Cas9* F_0_ (and therefore ♂ *w*^GDe^ F_0_) crosses. Positives are +*w* female F_2_s which should only occur through recombination. For ♀*Cas9* F_0_ crosses male F_2_s are considered. In each case, a Fisher’s exact test is performed with 13/1203 as the expected outcome (this includes recombination of the *white* allele in the -*Cas9* cross). Significance thresholds: * for <=0.05, ** for <=0.01, *** for <=0.001.

| Cas9 | Drive F<sub>1</sub> | Cas9 F<sub>0</sub> | Positives | Total Events | % | p-value | Threshold | Odds ratio |
| --- | --- | --- | --- | --- | --- | --- | --- | --- |
| -Cas9 | ♂ | N/A | 11 | 592 | 1.9% | 1.95E-01 | ns | 1.7 |
| *sds3* | ♂ | ♂ | 41 | 216 | 19.0% | 6.79E-22 | *** | 17.5 |
| *sds3* | ♂ | ♀ | 19 | 102 | 18.6% | 4.93E-13 | *** | 17.2 |
| *bgn* | ♂ | ♂ | 24 | 464 | 5.2% | 3.99E-06 | *** | 4.8 |
| *bgn* | ♂ | ♀ | 32 | 160 | 20.0% | 2.89E-19 | *** | 18.4 |
| *nup50* | ♂ | ♂ | 210 | 869 | 24.2% | 3.50E-57 | *** | 22.3 |
| *nup50* | ♂ | ♀ | 315 | 1387 | 22.7% | 1.02E-61 | *** | 21.0 |
| Li et al. *nup50* | ♂ | ♂ | 3 | 690 | 0.4% | 1.93E-01 | ns | 0.4 |
| Li et al. *nup50* | ♂ | ♀ | 3 | 688 | 0.4% | 1.94E-01 | ns | 0.4 |

**Table S16.**
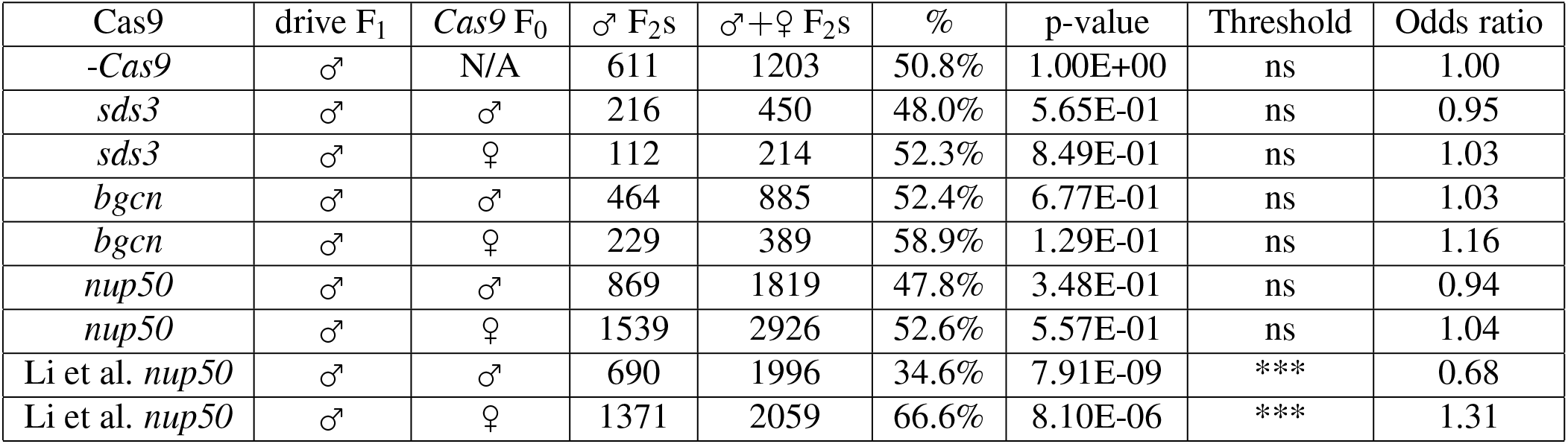
Meiotic drive significance test based on sex bias in F_2_ progeny. In each case, a Fisher’s exact test is performed with 611/1203 as the expected outcome. Significance thresholds: * for <=0.05, ** for <=0.01, *** for <=0.001.

## References

1. Alphey, L. S., Crisanti, A., Randazzo, F. F. & Akbari, O. S. Opinion: Standardizing the definition of gene drive. Proc. Natl. Acad. Sci. 117, 30864–30867, DOI: 10.1073/pnas.2020417117 (2020).

2. Burt, A. & Trivers, R. Genes in Conflict: The Biology of Selfish Genetic Elements, vol. 1 (Harvard University Press, 2006).

3. Esvelt, K. M., Smidler, A. L., Catteruccia, F. & Church, G. M. Concerning RNA-guided gene drives for the alteration of wild populations. eLife 3, 1–21, DOI: 10.7554/eLife.03401 (2014).

4. Champer, J., Buchman, A. & Akbari, O. S. Cheating evolution: engineering gene drives to manipulate the fate of wild populations. Nat. Rev. Genet. 17, 146–159, DOI: 10.1038/nrg.2015.34 (2016).

5. Jinek, M. et al. A Programmable Dual-RNA-Guided DNA Endonuclease in Adaptive Bacterial Immunity. Science 337, 816–821, DOI: 10.1126/science.1225829 (2012).

6. Oberhofer, G., Ivy, T. & Hay, B. A. Gene drive and resilience through renewal with next generation Cleave and Rescue selfish genetic elements. Proc. Natl. Acad. Sci. 117, 9013–9021, DOI: 10.1073/pnas.1921698117 (2020).

7. Oberhofer, G., Ivy, T. & Hay, B. A. Cleave and Rescue, a novel selfish genetic element and general strategy for gene drive. Proc. Natl. Acad. Sci. 116, 6250–6259, DOI: 10.1073/pnas.1816928116 (2019).

8. Champer, J. et al. A toxin-antidote CRISPR gene drive system for regional population modification. Nat. Commun. 11, 1082, DOI: 10.1038/s41467-020-14960-3 (2020).

9. Windbichler, N., Papathanos, P. A. & Crisanti, A. Targeting the X Chromosome during Spermatogenesis Induces Y Chromosome Transmission Ratio Distortion and Early Dominant Embryo Lethality in Anopheles gambiae. PLoS Genet. 4, e1000291, DOI: 10.1371/journal.pgen.1000291 (2008).

10. Galizi, R. et al. A synthetic sex ratio distortion system for the control of the human malaria mosquito. Nat. Commun. 5, 3977, DOI: 10.1038/ncomms4977 (2014).

11. Hickey, W. A. & Craig, G. B. J. Genetic distortion of sex ratio in a mosquito, Aedes aegypti. Genetics 53, 1177–1196 (1966).

12. Sweeny, T. L. & Barr, A. R. Sex Ratio Distortion Caused by Meiotic Drive in a Mosquito, Culex pipiens L. Genetics 88, 427–446 (1978).

13. Adolfi, A. et al. Efficient population modification gene-drive rescue system in the malaria mosquito Anopheles stephensi. Nat. Commun. 11, 5553, DOI: 10.1038/s41467-020-19426-0 (2020).

14. Gantz, V. M. et al. Highly efficient Cas9-mediated gene drive for population modification of the malaria vector mosquito Anopheles stephensi. Proc. Natl. Acad. Sci. 112, E6736–E6743, DOI: 10.1073/pnas.1521077112 (2015).

15. Pham, T. B. et al. Experimental population modification of the malaria vector mosquito, Anopheles stephensi. PLOS Genet. 15, e1008440, DOI: 10.1371/journal.pgen.1008440 (2019).

16. Kyrou, K. et al. A CRISPR–Cas9 gene drive targeting doublesex causes complete population suppression in caged Anopheles gambiae mosquitoes. Nat. Biotechnol. 36, 1062–1066, DOI: 10.1038/nbt.4245 (2018).

17. Hammond, A. et al. A CRISPR-Cas9 gene drive system targeting female reproduction in the malaria mosquito vector Anopheles gambiae. Nat. Biotechnol. 34, 78–83, DOI: 10.1038/nbt.3439 (2016).

18. Hammond, A. M. et al. The creation and selection of mutations resistant to a gene drive over multiple generations in the malaria mosquito. PLOS Genet. 13, e1007039, DOI: 10.1371/journal.pgen.1007039 (2017).

19. Carballar-Lejarazú, R. et al. Next-generation gene drive for population modification of the malaria vector mosquito, Anopheles gambiae. Proc. Natl. Acad. Sci. 117, 22805–22814, DOI: 10.1073/pnas.2010214117 (2020).

20. Champer, S. E. et al. Computational and experimental performance of CRISPR homing gene drive strategies with multiplexed gRNAs. Sci. Adv. 6, eaaz0525, DOI: 10.1126/sciadv.aaz0525 (2020).

21. Gantz, V. M. & Bier, E. The mutagenic chain reaction: A method for converting heterozygous to homozygous mutations. Science 348, 442–444, DOI: 10.1126/science.aaa5945 (2015).

22. Champer, J. et al. Reducing resistance allele formation in CRISPR gene drive. Proc. Natl. Acad. Sci. 115, 5522–5527, DOI: 10.1073/pnas.1720354115 (2018).

23. KaramiNejadRanjbar, M. et al. Consequences of resistance evolution in a Cas9-based sex conversion-suppression gene drive for insect pest management. Proc. Natl. Acad. Sci. 115, 6189–6194, DOI: 10.1073/pnas.1713825115 (2018).

24. Oberhofer, G., Ivy, T. & Hay, B. A. Behavior of homing endonuclease gene drives targeting genes required for viability or female fertility with multiplexed guide RNAs. Proc. Natl. Acad. Sci. 115, E9343–E9352, DOI: 10.1073/pnas.1805278115 (2018).

25. Champer, J. et al. Novel CRISPR/Cas9 gene drive constructs reveal insights into mechanisms of resistance allele formation and drive efficiency in genetically diverse populations. PLOS Genet. 13, e1006796, DOI: 10.1371/journal.pgen.1006796 (2017).

26. López Del Amo, V. et al. A transcomplementing gene drive provides a flexible platform for laboratory investigation and potential field deployment. Nat. Commun. 11, 352, DOI: 10.1038/s41467-019-13977-7 (2020).

27. Wu, B., Luo, L. & Gao, X. J. Cas9-triggered chain ablation of cas9 as a gene drive brake. Nat. Biotechnol. 34, 137–138, DOI: 10.1038/nbt.3444 (2016).

28. Xu, X.-r. S. et al. Active Genetic Neutralizing Elements for Halting or Deleting Gene Drives. Mol. Cell 80, 246–262, DOI: 10.1016/j.molcel.2020.09.003 (2020).

29. Kandul, N. P. et al. Assessment of a Split Homing Based Gene Drive for Efficient Knockout of Multiple Genes. G3; Genes|Genomes|Genetics 10, 827–837, DOI: 10.1534/g3.119.400985 (2020).

30. Champer, J. et al. Molecular safeguarding of CRISPR gene drive experiments. eLife 8, 1–10, DOI: 10.7554/eLife.41439 (2019).

31. Xu, X.-R. S., Gantz, V. M., Siomava, N. & Bier, E. CRISPR/Cas9 and active genetics-based trans-species replacement of the endogenous Drosophila kni-L2 CRM reveals unexpected complexity. eLife 6, 1–23, DOI: 10.7554/eLife.30281 (2017).

32. López Del Amo, V. et al. Small-Molecule Control of Super-Mendelian Inheritance in Gene Drives. Cell Reports 31, 107841, DOI: 10.1016/j.celrep.2020.107841 (2020).

33. Chae, D. et al. Chemical Controllable Gene Drive in Drosophila. ACS Synth. Biol. 9, 2362–2377, DOI: 10.1021/acssynbio.0c00117 (2020).

34. Champer, J. et al. A CRISPR homing gene drive targeting a haplolethal gene removes resistance alleles and successfully spreads through a cage population. Proc. Natl. Acad. Sci. 117, 24377–24383, DOI: 10.1073/pnas.2004373117 (2020).

35. Guichard, A. et al. Efficient allelic-drive in Drosophila. Nat. Commun. 10, 1640, DOI: 10.1038/s41467-019-09694-w (2019).

36. Li, M. et al. Development of a confinable gene drive system in the human disease vector Aedes aegypti. eLife 9, 1–40, DOI: 10.7554/eLife.51701 (2020).

37. Grunwald, H. A. et al. Super-Mendelian inheritance mediated by CRISPR–Cas9 in the female mouse germline. Nature 566, 105–109, DOI: 10.1038/s41586-019-0875-2 (2019).

38. Chan, Y.-S., Huen, D. S., Glauert, R., Whiteway, E. & Russell, S. Optimising Homing Endonuclease Gene Drive Performance in a Semi-Refractory Species: The Drosophila melanogaster Experience. PLoS ONE 8, e54130, DOI: 10.1371/journal.pone.0054130 (2013).

39. Simoni, A. et al. Development of synthetic selfish elements based on modular nucleases in Drosophila melanogaster. Nucleic Acids Res. 42, 7461–7472, DOI: 10.1093/nar/gku387 (2014).

40. Chan, Y.-S. et al. The Design and In Vivo Evaluation of Engineered I-OnuI-Based Enzymes for HEG Gene Drive. PLoS ONE 8, e74254, DOI: 10.1371/journal.pone.0074254 (2013).

41. Windbichler, N. et al. A synthetic homing endonuclease-based gene drive system in the human malaria mosquito. Nature 473, 212–215, DOI: 10.1038/nature09937 (2011).

42. Magnusson, K. et al. Transcription Regulation of Sex-Biased Genes during Ontogeny in the Malaria Vector Anopheles gambiae. PLoS ONE 6, e21572, DOI: 10.1371/journal.pone.0021572 (2011).

43. Bauer DuMont, V. L., Flores, H. A., Wright, M. H. & Aquadro, C. F. Recurrent Positive Selection at Bgcn, a Key Determinant of Germ Line Differentiation, Does Not Appear to be Driven by Simple Coevolution with Its Partner Protein Bam. Mol. Biol. Evol. 24, 182–191, DOI: 10.1093/molbev/msl141 (2007).

44. Anderson, M. A. In preparation. bioRxiv (2021).

45. Li, M. et al. Germline Cas9 expression yields highly efficient genome engineering in a major worldwide disease vector, Aedes aegypti. Proc. Natl. Acad. Sci. 114, E10540–E10549, DOI: 10.1073/pnas.1711538114 (2017).

46. Yusa, K., Zhou, L., Li, M. A., Bradley, A. & Craig, N. L. A hyperactive piggyBac transposase for mammalian applications. Proc. Natl. Acad. Sci. 108, 1531–1536, DOI: 10.1073/pnas.1008322108 (2011).

47. Coates, C. J., Jasinskiene, N., Miyashiro, L. & James, A. A. Mariner transposition and transformation of the yellow fever mosquito, Aedes aegypti. Proc. Natl. Acad. Sci. 95, 3748–3751, DOI: 10.1073/pnas.95.7.3748 (1998).

48. Matthews, B. J. et al. Improved reference genome of Aedes aegypti informs arbovirus vector control. Nature 563, 501–507, DOI: 10.1038/s41586-018-0692-z (2018).

49. Giraldo-Calderón, G. I. et al. VectorBase: an updated bioinformatics resource for invertebrate vectors and other organisms related with human diseases. Nucleic Acids Res. 43, D707–D713, DOI: 10.1093/nar/gku1117 (2015).

50. Pfitzner, C. et al. Progress Toward Zygotic and Germline Gene Drives in Mice. The CRISPR J. 3, 388–397, DOI: 10.1089/crispr.2020.0050 (2020).

51. Champer, J. et al. A CRISPR homing gene drive targeting a haplolethal gene removes resistance alleles and successfully spreads through a cage population. Proc. Natl. Acad. Sci. 117, 24377–24383, DOI: 10.1073/pnas.2004373117 (2020).

52. Terradas, G. et al. Inherently confinable split-drive systems in Drosophila. bioRxiv 2020.09.03.282079, DOI: 10.1101/2020.09.03.282079 (2020).

53. Hammond, A. et al. Regulation of gene drive expression increases invasive potential and mitigates resistance. bioRxiv 360339, DOI: 10.1101/360339 (2018).

54. Aryan, A. et al. Nix alone is sufficient to convert female Aedes aegypti into fertile males and myo-sex is needed for male flight. Proc. Natl. Acad. Sci. 117, 17702–17709, DOI: 10.1073/pnas.2001132117 (2020).

55. Kandul, N. P. et al. Transforming insect population control with precision guided sterile males with demonstration in flies. Nat. Commun. 10, 84, DOI: 10.1038/s41467-018-07964-7 (2019).

56. Adikusuma, F., Williams, N., Grutzner, F., Hughes, J. & Thomas, P. Targeted Deletion of an Entire Chromosome Using CRISPR/Cas9. Mol. Ther. 25, 1736–1738, DOI: 10.1016/j.ymthe.2017.05.021 (2017).

57. Zuo, E. et al. CRISPR/Cas9-mediated targeted chromosome elimination. Genome Biol. 18, 224, DOI: 10.1186/s13059-017-1354-4 (2017).

58. Fasulo, B. et al. A fly model establishes distinct mechanisms for synthetic CRISPR/Cas9 sex distorters. PLOS Genet. 16, e1008647, DOI: 10.1371/journal.pgen.1008647 (2020).

59. Xu, H. et al. Chromosome drives via CRISPR-Cas9 in yeast. Nat. Commun. 11, 4344, DOI: 10.1038/s41467-020-18222-0 (2020).

60. Ross, P. A., Endersby-Harshman, N. M. & Hoffmann, A. A. A comprehensive assessment of inbreeding and laboratory adaptation in Aedes aegypti mosquitoes. Evol. Appl. 12, 572–586, DOI: 10.1111/eva.12740 (2019).

61. Gloria-Soria, A., Soghigian, J., Kellner, D. & Powell, J. R. Genetic diversity of laboratory strains and implications for research: The case of Aedes aegypti. PLOS Neglected Trop. Dis. 13, e0007930, DOI: 10.1371/journal.pntd.0007930 (2019).

62. Noble, C. et al. Daisy-chain gene drives for the alteration of local populations. Proc. Natl. Acad. Sci. 116, 8275–8282, DOI: 10.1073/pnas.1716358116 (2019).

63. Oberhofer, G., Ivy, T. & Hay, B. A. 2-Locus Cleave and Rescue selfish elements harness a recombination rate-dependent generational clock for self limiting gene drive. bioRxiv 2020.07.09.196253, DOI: 10.1101/2020.07.09.196253 (2020).

